# Dominance shifts increase the likelihood of soft selective sweeps

**DOI:** 10.1101/2021.02.22.432386

**Authors:** Pavitra Muralidhar, Carl Veller

## Abstract

Genetic models of adaptation to a new environment have typically assumed that the alleles involved maintain a constant fitness dominance across the old and new environments. However, theories of dominance suggest that this should often not be the case. Instead, the alleles involved should frequently shift from recessive deleterious in the old environment to dominant beneficial in the new environment. Here, we study the consequences of these expected dominance shifts for the genetics of adaptation to a new environment. We find that dominance shifts increase the likelihood that adaptation occurs from the standing variation, and that multiple alleles from the standing variation are involved (a soft selective sweep). Furthermore, we find that expected dominance shifts increase the haplotypic diversity of selective sweeps, rendering soft sweeps more detectable in small genomic samples. In cases where an environmental change threatens the viability of the population, we show that expected dominance shifts of newly beneficial alleles increase the likelihood of evolutionary rescue and the number of alleles involved. Finally, we apply our results to a well-studied case of adaptation to a new environment: the evolution of pesticide resistance at the *Ace* locus in *Drosophila melanogaster*. We show that, under reasonable demographic assumptions, the expected dominance shift of resistant alleles causes soft sweeps to be the most frequent outcome in this case, with the primary source of these soft sweeps being the standing variation at the onset of pesticide use, rather than recurrent mutation thereafter.

## 1 Introduction

A primary concern of evolutionary genetics is to understand the genetic processes that underlie organisms’ adaptation to their environments. An important goal in this field is therefore to understand the nature of the genetic variation from which adaptation to a new environment typically occurs. When adaptation to a new environment is partly or wholly due to fixation of a newly beneficial allele at a given locus (a ‘selective sweep’), the question arises whether this fixation typically proceeds from a single initial copy of the beneficial allele (a ‘hard selective sweep’) or from multiple distinct copies that were possibly already segregating at the time of the environmental change (a ‘soft selective sweep’) (Hermisson and Pennings 2005; Pritchard et al. 2010; Messer and Petrov 2013). The relative frequency of hard versus soft sweeps has been the subject of much recent discussion [e.g., Messer and Petrov (2013); Jensen (2014); Schrider and Kern (2017); Hermisson and Pennings (2017); Harris et al. (2018); Garud et al. (2021)].

In the classic model of a selective sweep in response to a change of environment, a mutation that was neutral or deleterious before the environmental change becomes beneficial after the environmental change (Orr and Betancourt 2001). This simple model has been studied intensively from the perspective of mathematical population genetics [e.g., Hermisson and Pennings (2005); Pennings and Hermisson (2006); Pritchard et al. (2010); Messer and Petrov (2013); Hermisson and Pennings (2017); Stephan (2019)], and has served as the theoretical foundation for much empirical work [e.g., Barrett and Schluter (2008); Messer and Petrov (2013); Garud et al. (2015); Schrider and Kern (2017)].

An assumption that is usually invoked in both theoretical and empirical studies of this model is that the fitness dominance of the focal allele is invariant across the environmental change—that is, the allele’s dominance with respect to its deleterious effect in the old environment is equal to its dominance with respect to its beneficial effect in the new environment [an exception is Orr and Betancourt (2001), discussed below]. The reason for this assumption is convenience: it simplifies theoretical calculations and buys a degree of freedom in empirical studies. However, it also sidesteps a rich, century-old literature on physiological and evolutionary theories of allelic dominance.

Physiological theories of dominance provide mechanistic explanations for the observation that loss-of-function mutations are typically recessive with respect to the wild-type allele while gain-of-function mutations are typically dominant. De Vries and Bateson pioneered this literature in the late 19th century (Falk 2001), but its most famous representation is in the Wright-Kacser-Burns theory of metabolic dominance (Wright 1934; Kacser and Burns 1981; Keightley 1996), which, by explicitly modeling the chemistry of metabolic pathways, showed that their operation is intrinsically robust to single loss-of-function (or decrease-of-function) mutations. Physiological theories of dominance have generally focused on genes encoding enzymes or other products with ‘quasi-catalytic’ properties. However, empirical work has shown that the predictions of the physiological theories in fact hold across a much broader set of gene categories, suggestive of a need for more general explanations of patterns of dominance (Phadnis and Fry 2005; Agrawal and Whitlock 2011).

Such explanations can be found in evolutionary theories of dominance, which seek to explain why beneficial alleles tend to be dominant while deleterious alleles tend to be recessive. This literature begins in the 1920s with Fisher’s mathematical demonstration that modifiers of an allele’s dominance are under positive selection to increase its dominance when it is beneficial and to decrease its dominance when it is deleterious (Fisher 1928). While Fisher’s treatment was abstract, subsequent work—in many cases guided by physiological theories of dominance—has developed more mechanistically-explicit evolutionary theories of dominance, based on, for example, the dynamics of metabolic pathways [reviewed in Bourguet (1999)], models of optimal gene expression (Hurst and Randerson 2000), and multidimensional fitness landscapes (Manna et al. 2011).

There are two distinct scenarios under which a focal allele can transition from deleterious before an environmental change to beneficial after. In the first scenario, the same phenotype is primarily under selection before and after the environmental change, but the direction of selection on this phenotype changes. For example, an allele that reduces limb length would be deleterious in an environment in which long limbs are favored, but would become beneficial in a new environment where short limbs are advantageous [e.g., Donihue et al. (2018)]. In this scenario, the focal allele’s fitness dominance is governed by its dominance with respect to the selected phenotype; since the selected phenotype does not change across the environmental shift, nor would we expect the focal allele’s fitness dominance to change significantly.

In the second scenario, the environmental change corresponds to a change in the phenotype that is primarily under selection. In this case, the focal allele’s effect on heterozygotes’ fitness is modulated through different phenotypes before and after the environmental change, and so the allele’s fitness dominance is not expected to remain constant. In fact, the evolutionary and physiological theories of dominance predict that the dominance of the focal allele should shift from recessive when it is deleterious to dominant when it is beneficial.

For a concrete example of this second scenario, consider the *Ace* locus in insects, where recent adaptation has occurred in response to pesticide use, forming an important case study in the selective sweeps literature [e.g., Karasov et al. (2010); Garud et al. (2015)] and evolutionary genetics more broadly [e.g., Bourguet et al. (1997); Lenormand et al. (1999)]. *Ace* encodes the enzyme acetylcholinesterase, which catalyzes the breakdown of acetylcholine at the neuromuscular junction (Hoffmann et al. 1992; Fournier and Mutero 1994; Mutero et al. 1994; Bourguet and Raymond 1998). Organophosphate pesticides, introduced in the mid-twentieth century, inhibit acetylcholinesterase by targeting its binding site (Fournier et al. 1993; Fournier and Mutero 1994; Mutero et al. 1994; Shi et al. 2004). Mutations at the *Ace* locus that alter the shape of the binding site can confer resistance to pesticide binding (Menozzi et al. 2004; Shi et al. 2004), and therefore can be beneficial in environments where pesticides are used (Bourguet and Raymond 1998). However, the reconfigured enzymes are intrinsically less efficient at binding acetylcholine itself (Hoffmann et al. 1992; Fournier and Mutero 1994), rendering them deleterious in pesticide-free environments (Shi et al. 2004). Thus, in some geographic regions, these ‘resistant’ mutations were deleterious before the onset of pesticide use and beneficial after, conforming to the classic model described above. Moreover, the phenotype that was primarily under selection also changed, from intrinsic enzymatic efficiency before the onset of pesticide use to the ability to evade pesticide binding after. Consistent with the prediction of theories of dominance, the beneficial effect of resistant alleles in pesticide environments—stemming from their ability to evade pesticide binding—has been shown to be dominant across several insect species (Bourguet and Raymond 1998; Charlesworth 1998). The dominance of the deleterious effect of resistant alleles in pesticide-free environments—stemming from their reduced enzymatic efficiency—has been shown to be partially or fully recessive in at least two insect species (Labbé et al. 2014; Zhang et al. 2015); measurement of the enzymatic activity of resistant alleles in *Drosophila melanogaster* further suggests that they should be recessive deleterious in pesticide-free environments (Shi et al. 2004).

If, as we expect, adaptation to a new environment often involves a change in the phenotype primarily under selection, then the alleles involved in this adaptation likely often shift from recessive deleterious in the old environment to dominant beneficial in the new environment. This obviously holds major implications for the genetics of adaptation to new environments. Intuitively, if the alleles underlying adaptation to a new environment were recessive deleterious beforehand, they will tend to have been present in greater numbers in the standing variation at the time of the environmental change, increasing the chance that multiple alleles were involved in a subsequent selective sweep. That is, the pattern of dominance shifts predicted by the physiological and evolutionary theories is expected to increase the relative likelihood of soft versus hard selective sweeps, as well as the importance of alleles that were present in the standing variation at the time of the environmental change (versus those produced by mutation after the environmental change). Here, we carry out a quantitative investigation of the effect of these dominance shifts on the genetics of selective sweeps.

## 2 Methods

### The model

We study the classic model of a selective sweep in response to a change in the selective environment, adopting the framework set out by Hermisson and Pennings (2005). At a given locus, there are two alleles: the wild-type *A* and the mutant *a*. At a discrete point in time, *T*, there is a sudden environmental change. Prior to *T*, the mutant allele *a* was deleterious, with the relative fitnesses of genotypes *AA, Aa*, and *aa* being 1, 1 − *h*_*d*_*s*_*d*_, and 1 − *s*_*d*_, such that *h*_*d*_ is the fitness dominance of *a* prior to *T*. After *T, a* becomes beneficial, with the relative fitnesses of genotypes *AA, Aa*, and *aa* being 1, 1 + *h*_*b*_*s*_*b*_, and 1 + *s*_*b*_, such that *h*_*b*_ is the fitness dominance of *a* after *T*.

The population is of constant size *N* (= 10,000 in all simulations, unless otherwise stated), and evolves according to a Wright-Fisher process. Prior to *T*, the alleles *A* and *a* mutate to one another at a constant, symmetric rate *u* per replication. After *T*, there is no mutation, allowing us more precisely to study the likelihood and nature of adaptation from the standing variation (although we do later consider recurrent mutation after *T*).

### Definition of a soft sweep

Several definitions of a ‘soft sweep’ exist in the literature (Hermisson and Pennings 2017). These definitions can be partitioned according to two axes. First, some definitions consider the ancestry of the entire population of alleles after the sweep, while others consider the ancestry of only a sub-sample. Second, for a sweep from the standing variation to be called soft, some definitions require only that multiple alleles present at the time of the environmental shift have descendants upon completion of the sweep, while other definitions further require that those ancestral alleles have distinct mutational origins [this axis distinguishes ‘single-origin’ and ‘multiple-origin’ soft sweeps in the terminology of Hermisson and Pennings (2017)].

We primarily employ a definition that uses the first option along each of these two axes. By this definition, a sweep from the standing variation is soft if multiple copies of the allele that were present at the time of the environmental shift have descendants among the entire population of alleles upon completion of the sweep. We call this a ‘population’ definition of soft sweeps. However, some of our results relate directly to the empirical detectability of soft sweeps—see, e.g., ‘Measuring the haplotypic diversity of a sweep’ below. For these results, we employ a ‘sample’ definition of soft sweeps, using the second option along the two axes above. By this definition, a sweep is soft if, in a given sub-sample of alleles at the time of fixation, there are multiple mutational origins (and therefore multiple distinguishable haplotypes).

Note that the sample-based definition of a soft sweep is stricter than the population-based definition— fewer sweeps will be classified as soft under the sample definition. Thus, the population and sample definitions provide liberal and conservative predictions of the number of soft sweeps expected to occur, which could therefore be interpreted as upper and lower bounds.

### Simulation setup

We first characterize the mutation-selection-drift frequency distribution of *a* before *T* under various configurations of the parameters *u, s*_*d*_, and *h*_*d*_. For each configuration, we start in Hardy-Weinberg equilibrium at the focal locus, with the frequency of *a* equal to its large-population expectation (*u/*[*h*_*d*_*s*_*d*_] if 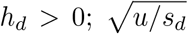 if *h*_*d*_ = 0). From this starting point, we allow the population to evolve for an initial burn-in period of 10^6^ generations, and thereafter record the frequency of each genotype every generation for 10^7^ generations. (In simulations where *N* = 100,000, owing to the greater computational expense, we recorded genotype frequencies for 3*×*10^6^ generations.) The distribution across generations of these genotype frequencies constitutes our empirical mutation-selection-drift distribution. In addition to recording the genotype frequencies in each generation, we record each distinct mutational origin of an *a* allele and the number of its descendant copies among *Aa* and *aa* genotypes.

We then study adaptation after *T*. For a given configuration of the parameters (*u, s*_*d*_, *h*_*d*_, *s*_*b*_, *h*_*b*_), we carry out 10,000 replicates of the following simulation: First, we randomly draw a set of starting genotype frequencies from the before-*T* mutation-selection-drift distribution for (*u, s*_*d*_, *h*_*d*_), estimated as described above. If one or more copies of *a* are present in this initial genotype configuration, we tag each separate copy, and track the descendants of each of these copies in every subsequent generation. The simulation ends when *a* fixes or goes extinct.

For each parameter configuration, we calculate the proportion of trials that result in each of the four possible outcomes: (i) *No standing variation*. No copies of *a* were present in the initial genotype configuration. (ii) *Failed sweep*. There was at least one copy of *a* present in the initial genotype configuration, but *a* subsequently went extinct. (iii) *Hard sweep. a* fixes, and all copies of *a* at the time of fixation descend from a single ancestral copy in the initial genotype configuration. (iv) *Soft sweep. a* fixes, and the copies of *a* at the time of fixation descend from more than one copy in the initial genotype configuration. Note that this analysis employs the ‘population definition’ of a soft sweep. In addition, in those trials in which a sweep occurred, we calculate the number of distinct mutational origins present among the swept alleles, and the frequencies of these distinct mutations. All simulations were run in SLiM 3 (Haller and Messer 2019).

### Measuring the haplotypic diversity of a sweep

When a sweep has occurred from alleles that were present in the standing variation, we are interested in the haplotypic diversity of the sweep. The haplotypic diversity of a sweep is interesting not just from a theoretical perspective, but also for a practical empirical reason: when a soft sweep is mutationally more diverse, we have a better chance of being able to recognize, in a finite sample of sequenced alleles, that a soft sweep has occurred. Therefore, in measuring the haplotypic diversity of sweeps, we shall aim to use metrics with practical relevance to the empirical assessment of selective sweeps. In relating these metrics to the empirical assessment of soft sweeps, we shall use the ‘sample definition’ of a soft sweep, that the alleles involved have multiple mutational origins.

Suppose that, in a trial in which *a* sweeps to fixation, the alleles present upon completion of the sweep derive from *m* distinct mutations before *T*. We record *m*. Let the frequency of the descendants of ancestral mutation *i* at the time of fixation be *p*_*i*_, *i* = 1, …, *m*. First, for various *n*, we measure the mutational (or ‘haplotypic’) diversity of the sweep by calculating the Gini-Simpson diversity index of order 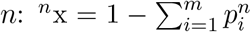 [modified from Jost (2006)]. Empirically, ^*n*^x is the probability that, in a random sample of *n* alleles taken from the population at the time of fixation, there are at least two distinct mutational origins and therefore at least two distinct haplotypes—i.e., ^*n*^x is the probability that a soft sweep from the standing variation can be detected in a random sample of *n* descendant alleles. Second, we record, for each value of *n*, the expected number of ancestral mutations represented in a random sample of *n* descendant alleles at the time of fixation, 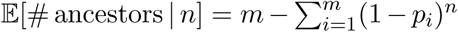. The formulae above are valid under the assumption that *n* ≪ *N*.

### Recurrent mutation

Selective sweeps can derive from copies of *a* present in the standing variation at the time of the environmental change, or from copies of *a* that appeared by mutation after the environmental change. To incorporate the possibility of recurrent mutation after the environmental change, we keep the symmetric *A* ↔ *a* mutation rate equal before and after the environmental change. Because *a* → *A* mutation after the environmental change generates a perpetual supply of *A* alleles, we say that *a* has ‘fixed’ (and a selective sweep has occurred) when it achieves a frequency equal to 0.99 times the mean of its large-population mutation-selection-drift distribution when beneficial.

Recurrent *A* → *a* mutation guarantees that a selective sweep will eventually occur after the environmental change. Therefore, we restrict our analysis of recurrent mutation to cases where a selective sweep occurs that involves alleles from the standing variation. We measure, in these cases, what proportion of *a* alleles present at the time of fixation derive from mutations that appeared after the environmental change versus mutations that were present in the standing variation at the time of the environmental change. Owing to the greater computational expense of these simulations, only 5,000 trials were run for some of the parameter configurations.

### Evolutionary rescue

The environmental change at time *T* could be such that the population would go extinct in the absence of the newly beneficial allele *a*. The question of a selective sweep of *a* is then one of evolutionary rescue. To study this situation requires abandoning our previous assumption of a constant population size, and explicitly modeling how the population shrinks or grows as a function of its genotypic composition.

We assume that the population is characterized by an intrinsic reproductive rate, *r*_0_. An interpretation of *r*_0_ is that, in a sexual population, and in the absence of any selective or ecological constraints, each individual would have 2(1 + *r*_0_) successful offspring on average; the population would then grow at rate *r*_0_ per generation. We scale ‘absolute’ fitnesses according to *r*_0_, such that an absolute fitness of *w* implies an expectation of 2(1 + *r*_0_)*w* successful offspring in the absence of any ecological constraints. The full population dynamics is then determined as follows. Suppose that, in generation *t*, the population is of size *N*_*t*_ and the average absolute fitness of its members is 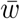. The ‘unconstrained’ population size in the next generation, 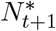, is a Poisson random variable with mean 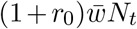; the actual population size in the next generation, *N*_*t*+1_, is the smaller of 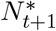 and the carrying capacity *K* = 10, 000. Once *N*_*t*+1_ is decided, the genotypic composition of generation *t* + 1 is determined by randomly drawing parental alleles from generation *t*, independently for each allele in generation *t* + 1, and with probabilities proportional to the fitnesses of the individuals carrying the alleles in generation *t*. Notice that, in the absence of the constraint that *N*_*t*+1_ *≤ K*, the ‘top down’ model described above would correspond to a simple ‘bottom up’ model where mating is random and the number of alleles contributed to generation *t* + 1 by an individual in generation *t* with absolute fitness *w* is a Poisson random variable with mean 2(1 + *r*_0_)*w*.

The general scenario for evolutionary rescue that we wish to model has the following key features: (i) Before *T*, the absolute fitness of the *AA* genotype, *w*_*AA*_, is such that (1 + *r*_0_)*w*_*AA*_ > 1, so that a population fixed (or nearly so) for *A* is held at its carrying capacity. (ii) After *T*, the absolute fitness of the *AA* genotype, 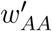, is such that 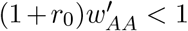, so that a population fixed for *A* would decline exponentially to extinction. (iii) After *T*, the absolute fitness of the *aa* genotype, 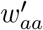, is such that 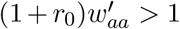, so that a population fixed for *a* would increase exponentially to the carrying capacity *K*. (iv) *a* is deleterious before *T* because of some impairment of basic function relative to *A*. After *T, a* confers resistance to whatever new selective force is threatening the population’s survival, but it still carries the cost of its impaired basic function. Therefore, 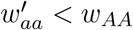.

To incorporate the above scenario into our model of selective sweeps, we assume that, before *T*, the absolute fitnesses of the genotypes *AA, Aa*, and *aa* are 1, 1 − *h*_1_*s*_1_, and 1 − *s*_1_ respectively, where *s*_1_ is the absolute fitness cost to *aa* individuals because of the impaired function of *a*, and *h*_1_ is the dominance of *a* relative to *A* with respect to this impaired function. Criterion (i) then simply requires that *r*_0_ > 0. After *T*, the absolute fitnesses of the genotypes *AA, Aa*, and *aa* are 1 − *s*_2_, (1 − *h*_2_*s*_2_)(1 − *h*_1_*s*_1_), and 1 − *s*_1_ respectively, where *s*_2_ is the absolute fitness cost to *AA* individuals because of *A*’s deleterious effect in the new environment (which *a* does not suffer), and *h*_2_ is the dominance of *A* relative to *a* with respect to this new deleterious effect. Criteria (ii) and (iii) require that (1 + *r*_0_)(1 − *s*_2_) < 1 and (1 + *r*_0_)(1 − *s*_1_) > 1, which in turn requires that *s*_2_ > *s*_1_, i.e., that the deleterious effect of *A* in the new environment is more severe than the deleterious effect of *a* in the old environment.

For each configuration of the parameters (*u, s*_1_, *h*_1_, *s*_2_, *h*_2_, *r*_0_), we carry out 10,000 replicates of the following simulation: First, we randomly draw a starting set of genotype frequencies from the before-*T* empirical mutation-selection-drift distribution corresponding for (*u, s*_1_, *h*_1_), estimated as described above. If one or more copies of *a* are present in this initial genotype configuration, we tag each separate copy, and track the descendants of each of these copies in every subsequent generation. We allow the population size and genotype frequencies to evolve according to the demographic model described above. The simulation is ended if the population either goes extinct (*N*_*t*_ = 0 for some *t*) or re-attains its carrying capacity (*N*_*t*_ = *K* for some *t*). In the latter case, the population has been rescued. Although the population size is initially expected to decline because the *A* allele is most common and (1 + *r*_0_)(1 − *s*_2_) < 1, we allow for chance fluctuations against selection in these early generations by only classifying a simulation run as an example of evolutionary rescue if *N*_*t*_ = *K* at least 10 generations after *T*. We do not allow for recurrent mutation after *T*.

In simulation runs where rescue is observed, we record: (i) The frequency of the *a* allele. This is of particular interest because, with fitnesses specified as in the model above, classical theories of dominance predict that there should often be heterozygote advantage at the focal locus in the new environment (see Results), in which case evolutionary rescue will involve only a partial sweep of *a*. (ii) The number of ancestral copies of *a* represented among descendant copies at the time of rescue. (iii) The haplotypic diversity among the population of *a* alleles at the time of rescue, the metrics for which are described above. (iv) The number of generations taken for rescue; i.e., the smallest *t* for which *N*_*t*_ = *K* (with the requirement that *t* > 10). (v) The minimum population size (i.e., how close the population came to extinction).

### Distribution of fitness effects

In the simulations described above, we assign fixed selection coefficients, *s*_*d*_ and *s*_*b*_, to the focal allele. To investigate the consequences of dominance shifts in a more general setting, we also ran simulations in which *s*_*d*_ and *s*_*b*_ were random variables, the realizations of which were drawn from empirically justified distributions of deleterious and beneficial fitness effects. Beneficial selection coefficients were drawn from an exponential distribution [e.g., Orr (2003); Eyre-Walker and Keightley (2007)] with mean 𝔼 [*s*_*b*_] = 0.01, a reasonable value that permits comparison with our fixed-effect simulations. Empirically, deleterious selection coefficients are often found to be well fit by a gamma distribution with shape parameter < 1 [e.g., Loewe et al. (2006)]. However, in this case, the mean frequency of the deleterious allele under mutation selection balance— 𝔼 [*u/*(*h*_*d*_*s*_*d*_)] if *h*_*d*_ > 0 and 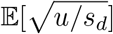 if *h*_*d*_ = 0—is undefined. Therefore, we instead drew deleterious selection coefficients from another empirically justified distribution, the lognormal distribution (Loewe and Charlesworth 2006; Kousathanas and Keightley 2013), calibrated to have the same mean and variance as a gamma distribution with shape parameter 0.5 [similar to estimates in *D. melanogaster* (Keightley and Eyre-Walker 2007; Schneider et al. 2011)] and mean 0.1 (permitting comparison with our fixed-effect simulations). Note that, while we have used simple, unimodal distributions for tractability and ease of comparison with our fixed-effect simulations, multimodal or mixed distributions of fitness effects can be preferable in some cases (Kousathanas and Keightley 2013; Bank et al. 2014; Johri et al. 2020).

In each replicate simulation, first *s*_*d*_ is drawn for the focal allele, independently across replicates, and the population dynamics proceed under the deleterious environment for 10^6^ generations (with bidirectional mutation at the focal locus, as before). Then *s*_*b*_ is drawn, independently across replicates and with respect to *s*_*d*_, and the population dynamics proceed under the beneficial environment until either fixation or loss of the focal allele. Owing to the greater computational expense of these simulations, 5,000 trials were run for each parameter setting.

## 3 Results

### Expected dominance shifts increase the likelihood of a selective sweep

A primary effect of the expected shift in dominance of the focal allele from recessive deleterious to dominant beneficial is to increase the probability that a selective sweep will occur after the environmental change. This result holds both when we treat the allele’s selection coefficients before and after the environmental change as fixed (Fig. 1A) or as random variables (Fig. S1). The increase in the probability of a sweep is especially pronounced when the rate of mutational supply, *θ* = 4*Nu*, is small (Fig. 1A).

**Figure 1:**
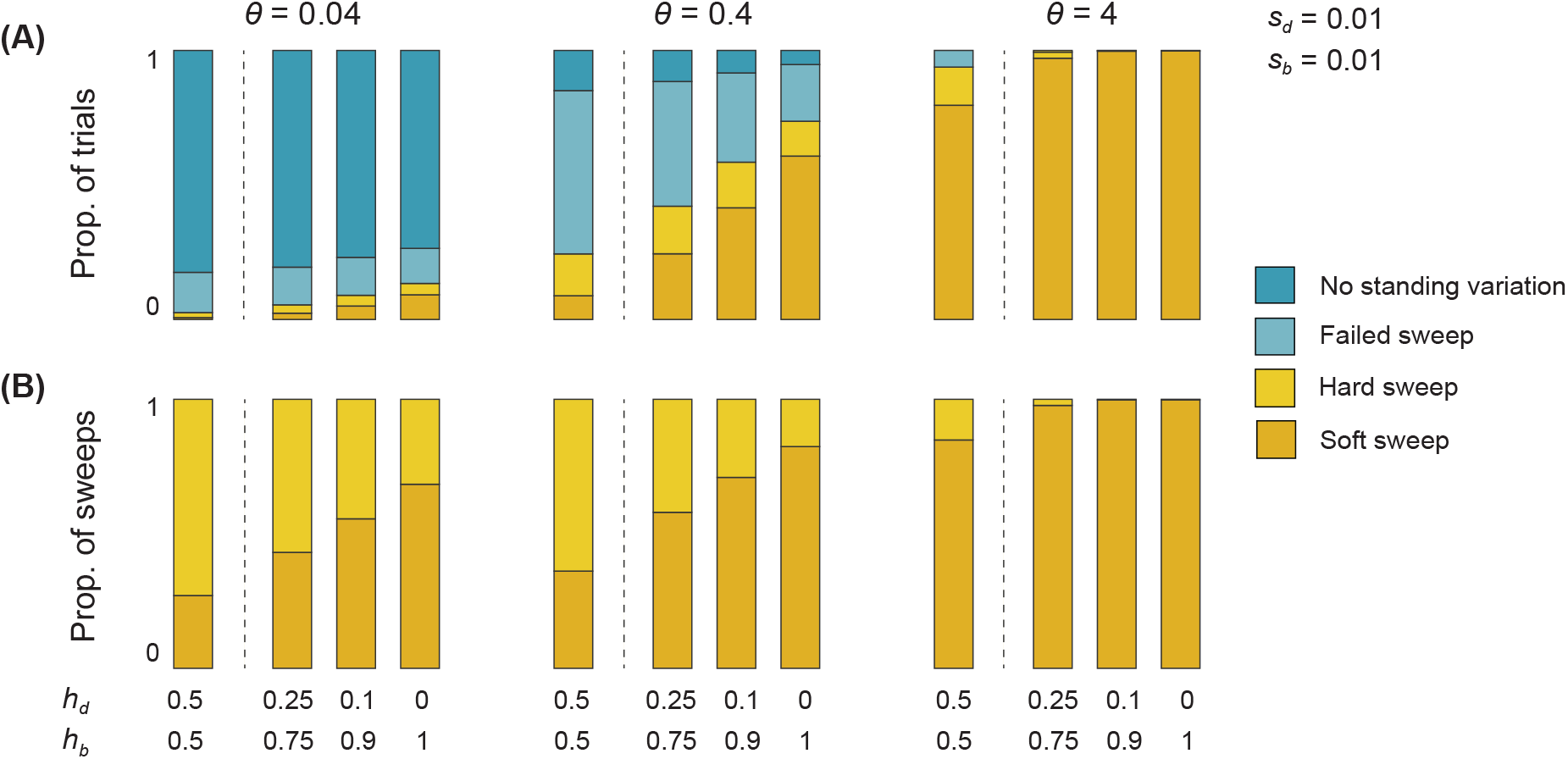
Expected dominance shifts increase the likelihood of a selective sweep and the proportion of selective sweeps that are soft. (A) The proportion of simulations that result in each of the four possible outcomes after the environmental change. The probability of a sweep (combined yellow and orange bars) increases with increasing severity of the focal allele’s dominance shift. This effect is proportionately largest when the rate of mutational supply, *θ* = 4*Nu*, is small. (B) Among trials in which a sweep does occur, the proportion in which multiple copies from the standing variation account for the descendant copies at the time of completion of the sweep. The proportion of such ‘soft sweeps’ increases with the severity of the dominance shift, with this effect again proportionately largest for small values of *θ*.

The reason for this increased probability of a sweep is straightforward. First, because the focal allele is recessive deleterious before the environmental shift, the mutation-selection-drift distribution is shifted towards there being more copies of the allele in the standing variation, relative to the case where the allele shows greater dominance in its deleterious effect. Second, the fact that the focal allele is dominant beneficial after the environmental change means that Haldane’s sieve does not hamper its initial chances of increasing in frequency and ultimately fixing. The first fact explains why the allele enjoys a higher probability of sweeping to fixation than a comparable allele that is dominant both before and after the environmental change; the second fact explains the same outcome with respect to an allele that is recessive both before and after the environmental change (Fig. S3).

### Expected dominance shifts increase the likelihood of soft versus hard sweeps

A dominance shift of the focal allele also increases the probability, conditional on a selective sweep occurring, that the sweep will derive from multiple copies of the allele that were present in the standing variation (a soft sweep by our ‘population definition’) relative to just one (a hard sweep). Again, this result holds for both fixed and random selection coefficients (Figs. 1B, S1).

This effect is clearly driven by the influence of dominance on the mutation-selection-drift distribution before the environmental change: when the allele is recessive deleterious before the environmental change, there are likely to be more copies present in the standing variation at the time of the environmental change, and so it is more likely that multiple copies will be involved in a subsequent sweep. The effect of dominance shifts in increasing the relative likelihood of soft versus hard selective sweeps is especially noticeable for small values of *θ* (Fig. 1B).

Importantly, for a given value of *θ*, the expected number of copies of the allele that were present as standing variation at the time of the environmental change differs conditional on a hard versus a soft sweep subsequently occurring (Fig. S2) (Hermisson and Pennings 2017). In other words, hard and soft sweeps tend to derive from different parts of the unconditional mutation-selection-drift distribution, underscoring the point that the mutation-selection-drift distribution of *a* before the environmental change—rather than just the mean of this distribution—must be considered to understand the population genetics of subsequent adaptation (Hermisson and Pennings 2017).

There has recently been much debate about the size of the parameter space under which soft sweeps prevail over hard sweeps (Messer and Petrov 2013; Jensen 2014; Hermisson and Pennings 2017; Harris et al. 2018; Garud et al. 2021). A corollary of the results above is that this parameter space is substantially expanded by the shifts in dominance predicted by the physiological and evolutionary theories of dominance.

Two parameters have been considered particularly relevant for the relative likelihood of hard versus soft sweeps: the rate of mutational supply of the focal allele (*θ*), and the ratio of the focal allele’s beneficial effect after the environmental change to its deleterious effect beforehand (*s*_*b*_*/s*_*d*_). On the first, soft sweeps are relatively unlikely for small values of *θ* (Hermisson and Pennings 2017). However, with dominance shifts of the focal allele, soft sweeps can be relatively likely for values as low as *θ* ∼ 0.01 (Fig. 1, left; Fig. S4). For higher values of *θ*, where soft sweeps predominate over hard sweeps even if the focal allele’s dominance is constant across the environmental change, dominance shifts have a relatively small effect on the likelihood of soft versus hard sweeps (Fig. 1, right). On the second, higher values of *s*_*b*_*/s*_*d*_ clearly make soft sweeps from the standing variation more likely, but there is uncertainty about the precise values of *s*_*b*_*/s*_*d*_ for which soft sweeps are expected to predominate (Jensen 2014; Hermisson and Pennings 2017). We find that dominance shifts drastically reduce the *s*_*b*_*/s*_*d*_ values required for soft sweeps to be likely—indeed, soft sweeps can predominate over hard sweeps for values as low as *s*_*b*_*/s*_*d*_ = 0.1 in certain parameter regimes (Fig. S4), and can predominate more generally when *s*_*b*_*/s*_*d*_ = 1 (Fig. 1).

These effects are strongest when the shift in dominance of the focal allele is complete, but more modest shifts in dominance can also cause soft sweeps to have an appreciable likelihood relative to hard sweeps for low values of *θ* and *s*_*b*_*/s*_*d*_ (Figs. 1, S5). Note that, when soft sweeps are defined according to the stricter ‘sample definition’, they become less likely. However, dominance shifts nevertheless increase the probability of soft sweeps according to this definition, and they are still possible for *θ* values as low as ∼0.01 (Fig. S6).

### Expected dominance shifts lead to greater haplotypic diversity within soft sweeps

Expected dominance shifts also cause selective sweeps, when they do occur, to be mutationally (and therefore haplotypically) more diverse, according to the Gini-Simpson diversity index for various orders *n* (Fig. 2A; see Methods). A practical consequence is that dominance shifts cause soft selective sweeps to be more detectable in small genomic samples, relative to the case where the focal allele’s dominance remains constant across the environmental change. Similarly, under dominance shifts, a greater number of mutational lineages (and therefore haplotypes) are expected to be present in a sample of alleles taken at the time of completion of the sweep (Fig. 2B).

**Figure 2:**
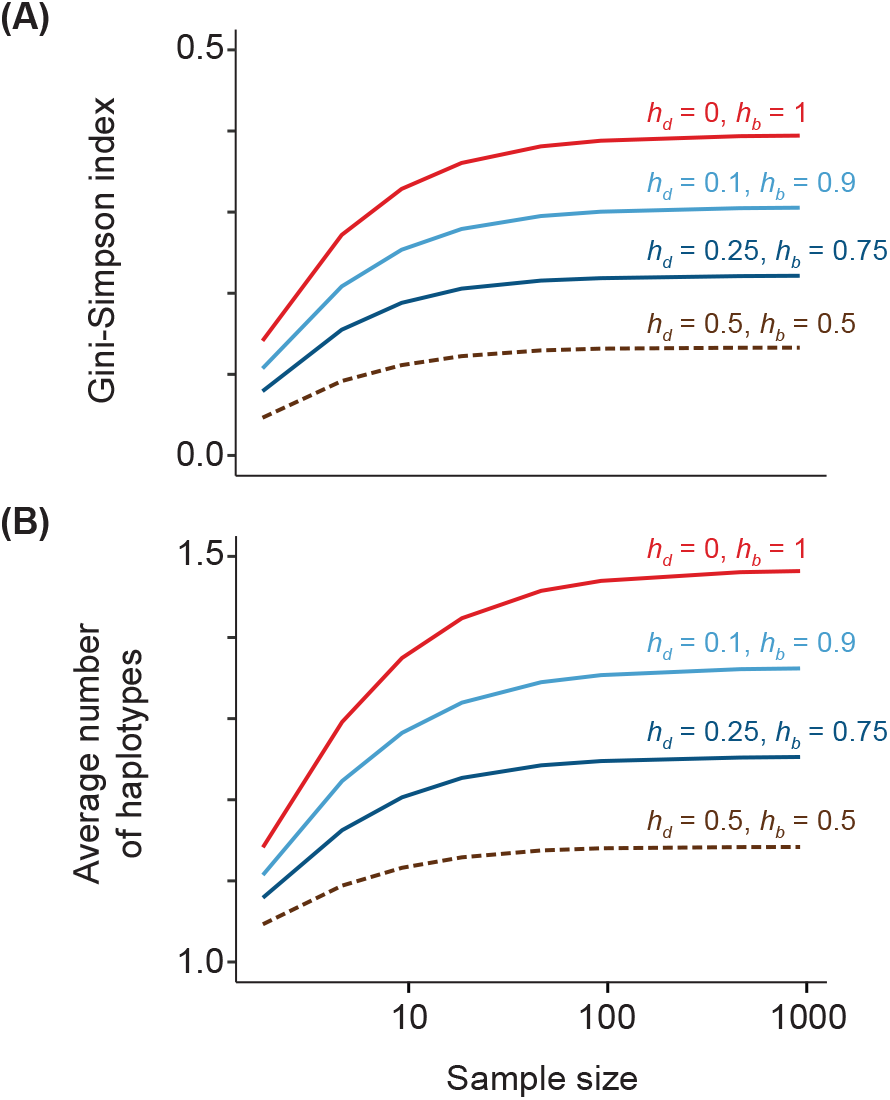
Expected dominance shifts increase the allelic diversity—the ‘softness’—of selective sweeps. Compared to the case where the focal allele shows constant additive fitness dominance before and after the environmental change (brown dotted lines), dominance shifts increase (A) the Gini-Simpson diversity index of alleles involved in the sweep, and (B) the expected number of alleles involved in the sweep. Consequently, dominance shifts cause soft sweeps to be detectable more often in samples of alleles taken upon completion of the sweep. Parameters: *s*_*d*_ = 0.01, *s*_*b*_ = 0.01.

Thus, for the parameters considered in Fig. 2, and in a sample of 10 alleles taken upon completion of a sweep, a soft sweep would be detected only 11% of the time if the allele maintained a constant additive dominance before and after the environmental change (the expected number of haplotypes present in the sample is 1.11 in this case), but would be detected 33% of the time if the allele instead underwent the dominance shift predicted by the evolutionary and physiological theories of dominance (and the expected number of haplotypes in the sample would rise to 1.37).

### The importance of population size, aside from *θ*

In the simulations described above, we have assumed a population size of *N* = 10,000, and have varied the mutation rate *u*, rather than *N*, to study the effect of the rate of mutational supply, *θ* = 4*Nu*. This choice was made for computational efficiency. In most cases, it is not expected to have a substantial impact on our results. To see this, first, consider the case where the focal allele shows substantial dominance before and after the environmental change (*h*_*d*_, *h*_*b*_ ≫ 0). In this case, the distribution of the number of copies of the allele in the standing variation at the time of the environmental change is controlled predominantly by *θ*, and will not shift much for different values of *N* and *u* that result in the same value of *θ* (for example, the mean of the mutation-selection distribution is 2*N × u/*[*h*_*d*_*s*_*d*_] = *θ/*[2*h*_*d*_*s*_*d*_]). Once the allele becomes beneficial, it is the number of copies present in the standing variation, rather than the fraction of the population that they constitute, that matters for the probability of a sweep (and the probability that a sweep will be soft, by the population definition). Therefore, in this case, *N* will not substantially affect the probability of a sweep, and of a soft sweep, except through its effect on *θ* (Fig. S7A,C).

Now consider the case where the allele is recessive both before and after the environmental change (*h*_*d*_, *h*_*b*_ = 0). In this case, the distribution of the number of copies of the allele present in the standing variation at the time of the environmental change does depend on *N* independently of *N* ‘s effect on *θ* (for example, the mean of the mutation-selection distribution is approximately 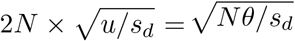. However, once the allele becomes beneficial, it is not the number of copies of the allele that matters for the probability of a sweep (and of a soft sweep), but rather the number of individuals homozygous for the allele. The distribution of the number of homozygotes is determined by *θ* (for example, its mean is 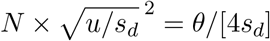), and so, again, the probability of a sweep (and of a soft sweep) will not substantially depend on *N* apart from *N* ‘s influence on *θ* (Fig. S7A,C).

Finally, consider the case where the allele undergoes the expected shift in dominance, from recessive to dominant (*h*_*d*_ = 0, *h*_*b*_ ≫ 0). In this case, the distribution of the number of copies of the allele present at the time of the environmental change does depend on *N* independently of *N* ‘s effect on *θ*—scaling approximately with 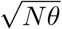, as described above—and the probability of a sweep (and a soft sweep) occurring once the allele becomes beneficial does depend on the number of copies of the allele present (rather than the number of homozygotes). Therefore, in this case, population size itself influences the probability of a sweep (and of a soft sweep), independently of its effect on the mutational supply *θ*. In particular, holding *θ* constant, sweeps are more likely, (and more likely to be soft, in larger populations (Fig. S7A,C).

The observations above pertain to soft sweeps defined according to the weaker, population-based definition. However, they also hold for the stricter sample-based definition. Thus, when the focal allele shows substantial dominance before and after the environmental change, the relative probabilities of a multi- versus a single-haplotype sweep do not depend on *N*, if *θ* is held constant (Fig. S7B,D). However, if there is a strong dominance shift of the focal allele from recessive deleterious to dominant beneficial, then a sweep is substantially more likely to involve multiple haplotypes when *N* is larger, holding *θ* constant (Fig. S7B,D).

Thus, consideration of strong dominance shifts reveals an intriguing exception to the rule that population size influences the genetics of selective sweeps only through its effect on the mutational supply.

### Expected dominance shifts decrease the importance of recurrent mutation versus standing variation for soft selective sweeps

Selective sweeps can derive from alleles that were present in the standing variation at the time of the environmental change, or from alleles that appeared by mutation after the environmental change (Pennings and Hermisson 2006; Hermisson and Pennings 2017). Recurrent mutation after the environmental change guarantees that a sweep will eventually occur, which narrows our interest to two questions.

First, how often is recurrent mutation necessary for a selective sweep to occur? This is equivalent to asking how often a selective sweep occurs in the absence of recurrent mutation, using alleles from the standing variation alone. We have addressed this question above, and have shown that dominance shifts substantially increase the probability that a selective sweep—and a soft sweep in particular—will occur from alleles that were present in the standing variation at the time of the environmental change. Second, in cases where a selective sweep occurs and alleles from the standing variation are involved, how large a role do mutations that occurred after the environmental change play in the sweep?

To answer this question, we incorporate recurrent mutation into our simulation setup (see Methods). Previous theory has suggested that recurrent mutation should often play a leading role in selective sweeps, relative to the standing variation (Pennings and Hermisson 2006; Hermisson and Pennings 2017). We find that a dominance shift of the focal allele increases the importance of the standing variation as a source of alleles in selective sweeps (Fig. 3). The reason is straightforward. Relative to the case where the allele is dominant (or semi-dominant) before and after the environmental change, a dominance shift of the focal allele does not alter the nature of selection acting on copies of it produced by recurrent mutation after the environmental change—in both scenarios, such copies arise at rate proportional to *θ* and are dominant beneficial. However, the dominance shift does increase the number of copies of the allele present in the standing variation at the time of the environmental change, improving the prospects of the standing variation as an allelic source of a subsequent sweep. Thus, the expected dominance shift increases the importance of the standing variation relative to recurrent mutation.

**Figure 3:**
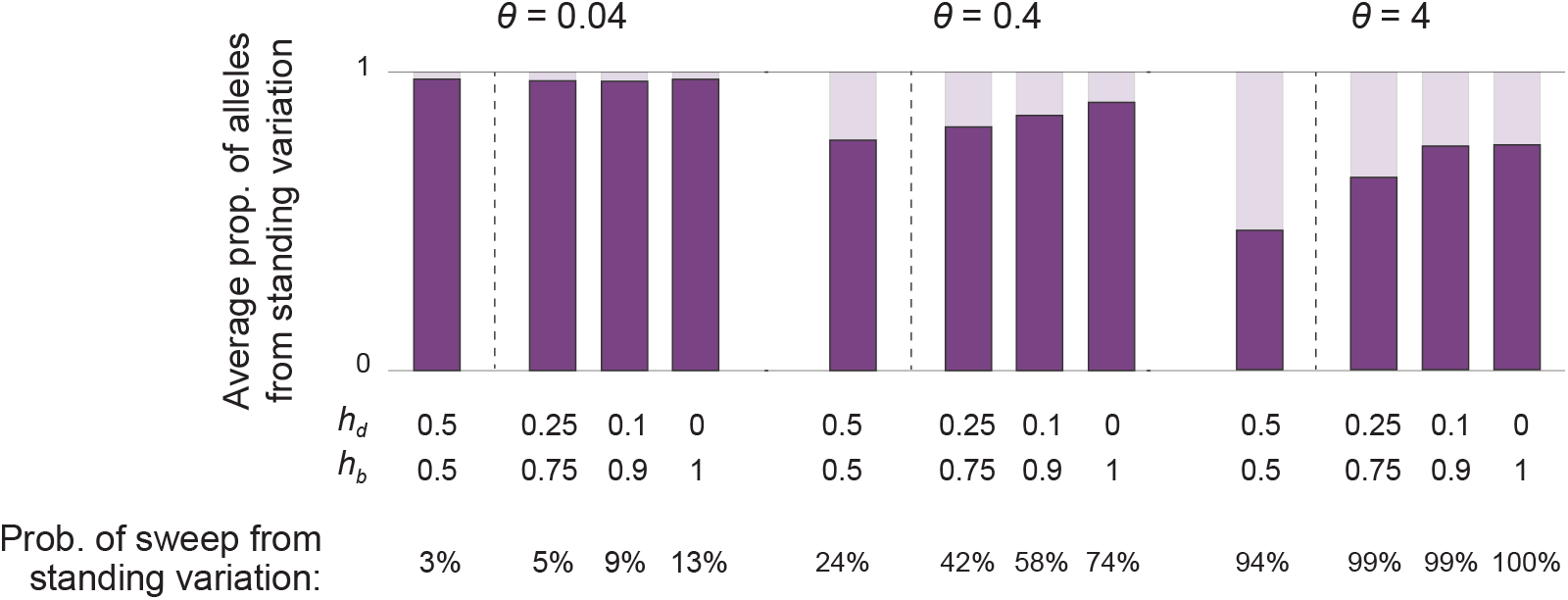
In selective sweeps partly sourced from the standing variation at the time of the environmental change, dominance shifts increase the representation of alleles from the standing variation relative to those produced by mutation after the environmental change. When *θ* is small (left panel), successful sweeps tend to be dominated by alleles from the standing variation, regardless of whether a dominance shift occurs (conditional on these sweeps involving at least one allele from the standing variation, the probabilities of which are displayed at the bottom of the figure). In contrast, when *θ* is large (right panel), mutations arising after the environmental change have more opportunity to ‘compete’ with alleles from the standing variation before the sweep is completed, and dominance shifts strongly increase the representation of alleles from the standing variation in successful sweeps. Parameters: *s*_*d*_ = 0.01, *s*_*b*_ = 0.01.

This effect of dominance shifts is most noticeable for high values of *θ* (e.g., *θ* ∼ 1), where the large supply of new mutations immediately after the environmental change increases the chance that these new alleles will be incorporated into a successful sweep. In contrast, for small values of *θ* (e.g., *θ* ∼ 0.01), the mutational supply of alleles immediately after the environmental change is small, so that sweeps—when they do occur—mostly involve alleles from the standing variation irrespective of their dominance before and after the environmental change.

### Expected dominance shifts increase the likelihood of evolutionary rescue

We have thus far assumed a constant population size, both before and after the environmental change. However, in many cases of interest, the relative benefit enjoyed by the mutant allele *a* after the environmental change will be due in part to a reduced absolute fitness of the wild-type allele *A* in the new environment, such that a population fixed for *A* would go extinct. In such cases, a selective sweep of *a* might be required for the population to recover in size (i.e., to be ‘rescued’).

Incorporating into our model the dependence of population size on mean absolute fitness (see Methods), we find that a dominance shift of *a* increases the probability of evolutionary rescue, relative to the case where the dominance of *a* remains constant across the environmental change (Fig. 4). Moreover, in cases where rescue does occur, the dominance shift of *a* reduces the severity of the bottleneck suffered by the population, and allows the population to re-attain its prior size more rapidly (Fig. 4).

**Figure 4:**
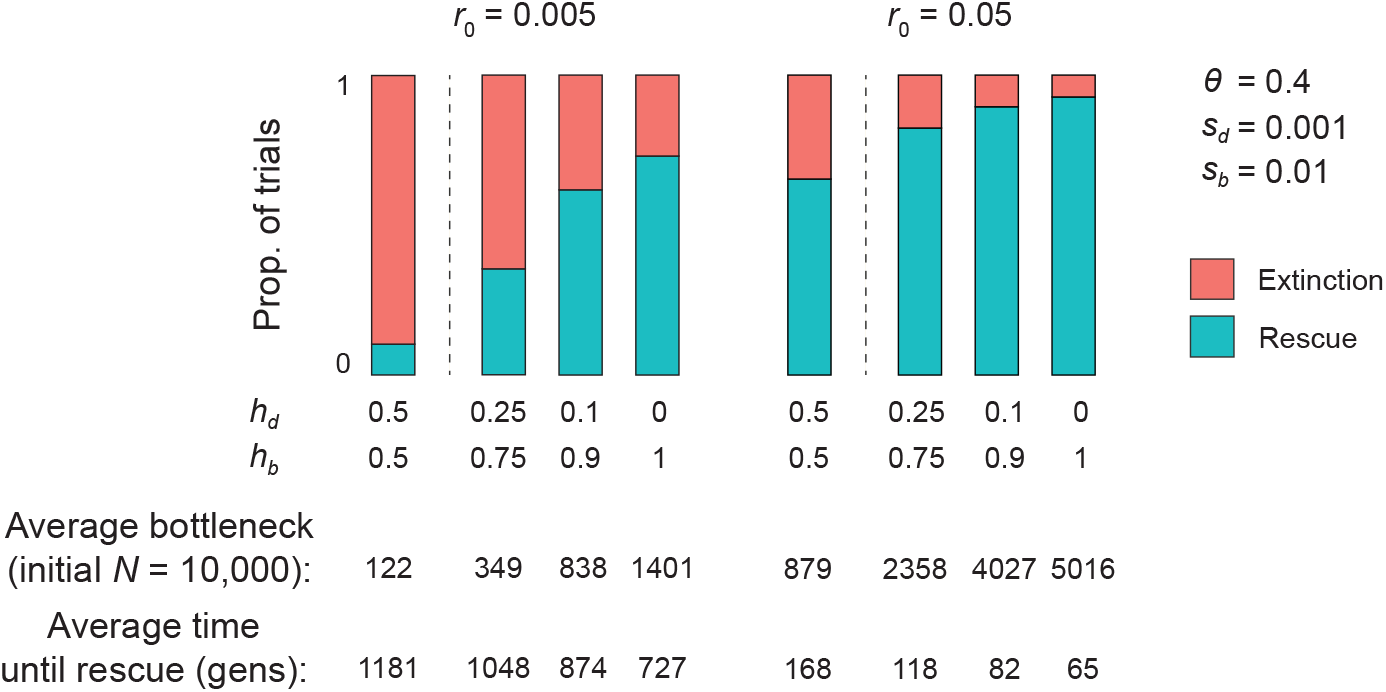
Expected dominance shifts increase the likelihood of evolutionary rescue in response to an environmental change that threatens the viability of the population. In cases where rescue does occur, it is more rapid, and the population size bottleneck less severe, when the rescuing allele undergoes a dominance shift. The likelihood of evolutionary rescue is especially increased by dominance shifts, relative to the case of constant dominance, when the population has a low intrinsic growth rate, *r*_0_ (left panel). However, dominance shifts especially increase the speed of rescue, and make less severe the population size bottleneck, when *r*_0_ is greater (right panel).

An interesting implication of expected dominance shifts in our model of evolutionary rescue is that heterozygote advantage after the environmental shift should be a common outcome. In this model, *a* is associated with a relatively small absolute fitness cost in both the pre- and post-change environments, while *A* is associated with a relatively large absolute fitness cost in the post-change environment. Therefore, *a* is relatively disadvantageous before the environmental change, but relatively advantageous after (making possible evolutionary rescue). When *a* is recessive with respect to the absolute fitness disadvantage it induces before the environmental change, but dominant with respect to the absolute fitness disadvantage induced by *A* after the environmental change, then *Aa* heterozygotes suffer neither of the absolute fitness reductions induced by the two alleles in the post-change environment, and therefore enjoy the highest relative fitness. In such cases, when evolutionary rescue does occur, it is expected to proceed via a partial sweep of *a*, resulting in a balanced polymorphism at the focal locus.

### Case study: Adaptation at the *Ace* locus

To demonstrate the empirical relevance of expected dominance shifts for selective sweeps, we consider their importance for a well-studied case of adaptation to a new environment in insects: adaptation at the *Ace* locus in response to pesticide use.

Karasov et al. (2010) collected sequence data at the *Ace* locus from pesticide-resistant and pesticide-sensitive strains of *Drosophila melanogaster*. Comparing the sequences of resistant alleles, they inferred multiple haplotypes, i.e., a soft selective sweep by our ‘sample definition’. Since the point mutations that confer resistance are known in this case (Menozzi et al. 2004), Karasov et al. (2010) were able to calibrate the standard selective sweeps model using the point mutation rate of *D. melanogaster*, the species’ traditionally quoted effective population size (*N*_*e*_ ∼ 10^6^), and known fitness parameters for resistant alleles at the *Ace* locus, but assuming the dominance of resistant alleles to be constant before and after the onset of pesticide use. Karasov et al. (2010) found that, under these parameters: (i) a sweep involving the standing variation would have been unlikely; (ii) a sweep seeded by mutation after the onset of pesticide use would likely have been hard.

To reconcile the selective sweeps model with their observation of a soft sweep at the *Ace* locus, Karasov et al. (2010) proposed that the relevant effective population size for recent adaptation (such as at the *Ace* locus) would not be based on the long-term demography of the species (as the traditional effective population size is), but would instead depend on more recent demography, owing to the short timescale over which the relevant adaptation has occurred. Because *D. melanogaster* has undergone a recent population expansion (Thornton and Andolfatto 2006), an effective population size based on recent demography would be substantially larger than the long-term effective population size. Substituting into the selective sweeps model an effective population size two orders of magnitude larger than the traditionally quoted quantity, Karasov et al. (2010) found that a multi-haplotype soft sweep—as observed in their data—would be the expected outcome.

Our results suggest another factor that would help to reconcile the surprisingly high haplotypic diversity of resistant alleles at the *Ace* locus with the classic model of a selective sweep: a dominance shift of resistant alleles, from recessive deleterious in pesticide-free environments to dominant beneficial in environments of pesticide use. Indeed, empirical evidence suggests that resistant alleles at *Ace* in insects exhibit this pattern of fitness dominances in the two environments (Bourguet and Raymond 1998; Charlesworth 1998; Zhang et al. 2015). To understand what effect such dominance shifts might have on expected diversity among resistant alleles, we consider a single-locus model, employing the same mutation and fitness parameters as Karasov et al. (2010), and we estimate the likelihood of a sweep, the likelihood that the sweep is soft (by both the population and sample definitions), and the expected haplotypic diversity within a sweep, for various degrees of dominance shift [including no shift, as considered by Karasov et al. (2010)] and for various effective population sizes.

First, we study the case of *θ* = 0.04, a value that corresponds approximately to the traditional value of the effective population size in *D. melanogaster* (*N*_*e*_ ∼ 10^6^). First, we find that the probability of adaptation from the standing variation is extremely small when there is no dominance shift of the resistant allele (*h*_*d*_ = *h*_*b*_ = 0.5) (Fig. 5A), consistent with Karasov et al. (2010). In addition, in the rare cases where a sweep does occur from the standing variation, the sweep usually involves only one copy of the resistant allele—and almost always only one haplotype (Fig. 5B,C). Under a full dominance shift of the resistant allele (*h*_*d*_ = 0; *h*_*b*_ = 1), the probability of adaptation from the standing variation increases substantially, but remains small, when *θ* = 0.04. When a sweep does occur, it now more often uses multiple copies of the resistant allele from the standing variation (Fig. 5B), but still typically involves only one haplotype (Fig. 5C). Thus, in this case, sweeps would typically be soft by the population definition, but these soft sweeps would seldom be detectable in genomic data.

**Figure 5:**
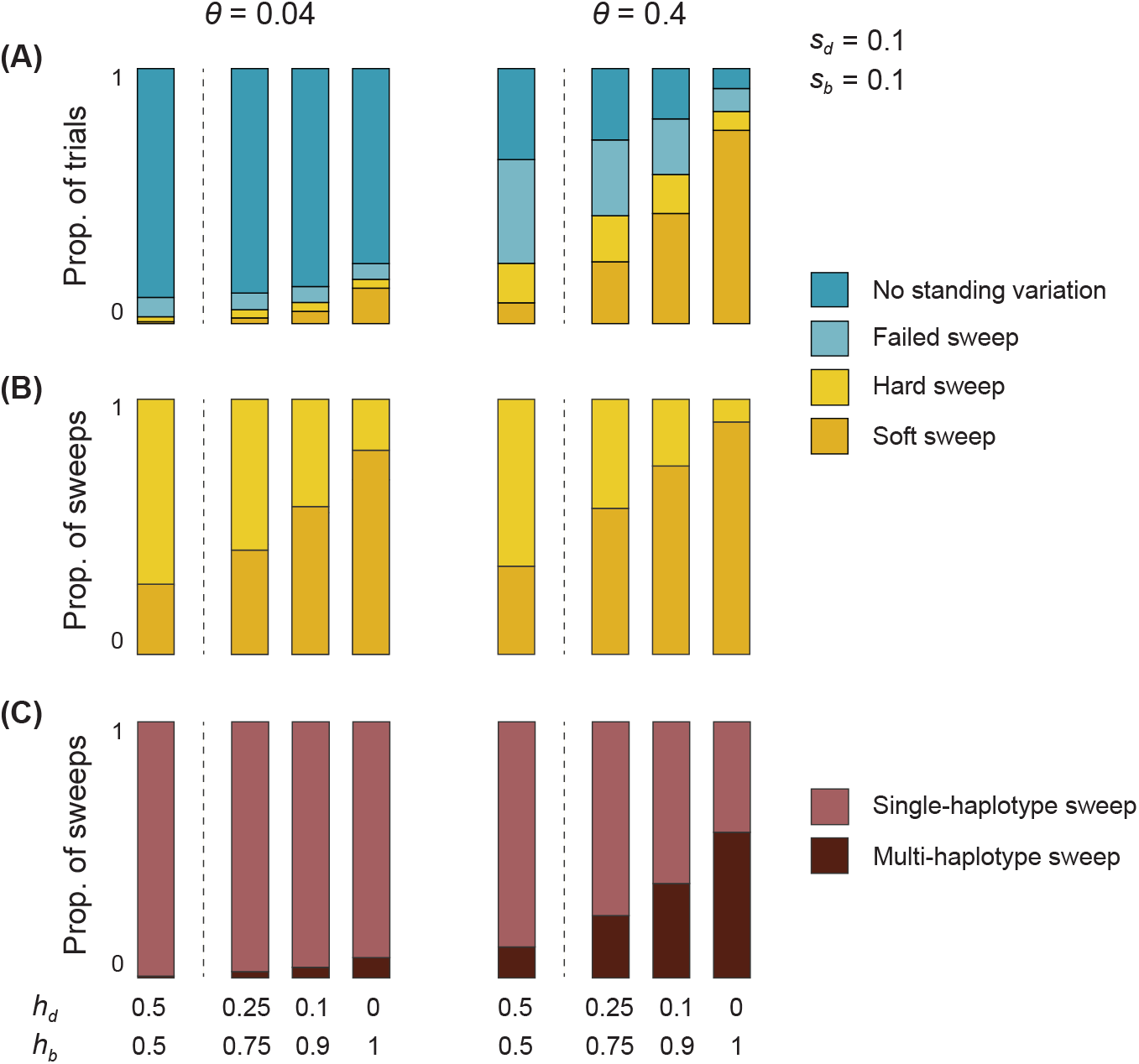
In response to the use of organophosphate pesticides, a dominance shift of resistant alleles at the *Ace* locus increases (A) the likelihood of a sweep from the standing variation, (B) the likelihood that a sweep from the standing variation is soft, and (C) the likelihood that multiple distinct haplotypes are involved in a sweep. These probabilities are displayed for values of *θ* corresponding to the long-term effective population size of *Drosophila melanogaster* (left panels) and, following the logic of Karasov et al. (2010), an increased estimate of the relevant effective population size based on more recent demography of the species (right panels). With this modestly revised estimate of the relevant effective population size, the soft sweep that Karasov et al. (2010) inferred from sequence data would be the expected outcome under a strong dominance shift of resistant alleles; moreover, alleles from the standing variation alone would usually be sufficient to furnish an empirically detectable soft sweep.

We now study the case of *θ* = 0.4, corresponding to a ten-fold higher effective population size than the traditional quantity for *D. melanogaster*, but a ten-fold lower value than that posited by Karasov et al. (2010) to explain the allelic diversity they observed at *Ace*. When the resistant allele maintains a constant additive dominance across the pre- and post-pesticide-use environments, adaptation from the standing variation remains unlikely in this case (Fig. 5A). Furthermore, when sweeps do occur, they typically make use of only one allele, and one haplotype, from the standing variation (Fig. 5B,C). However, when the resistant allele undergoes a full dominance shift, adaptation from the standing variation becomes the expected outcome (Fig. 5A). Now, when a sweep does occur, it almost always makes use of multiple copies of the allele from the standing variation (Fig. 5B; a soft sweep by the population definition), and more often than not involves multiple haplotypes (Fig. 5B), rendering the soft sweep empirically detectable in genomic data.

In the simulations described above, rather than fixing the mutation rate and varying the effective population size to simulate different values of *θ*, for computational efficiency we have instead fixed the effective population size at *N*_*e*_ = 10^4^ and varied the mutation rate. As discussed earlier, whereas these two procedures should ordinarily yield similar results, this is not so if the focal allele undergoes a substantial dominance shift: in this case, sweeps—and soft sweeps in particular—are more likely in larger populations, holding *θ* constant (Fig. S7). In our simulations of the *Ace* locus in *Drosophila*, if we consider different effective population sizes within a computationally feasible range (*N*_*e*_ ∈ *{*10^3^, 10^4^, 10^5^*}*) while holding *θ* fixed at 0.4, we find that the expected dominance shift of the resistant allele increases the probability of a sweep, and of a soft sweep, more drastically at larger population sizes, relative to the case where the allele does not undergo a dominance shift (Fig. S7C,D). Although realistic effective population sizes for *Drosophila* (*N*_*e*_ ≥ 10^6^) cannot feasibly be simulated with our setup, we can get a rough picture of what the results would look like at these population sizes by extrapolating the patterns we have observed across computationally feasible population sizes (Fig. S10). This exercise suggests that our estimates reported above of the increased likelihood of soft selective sweeps at the Ace locus induced by dominance shifts (and thus also the more modest degree to which the relevant effective population size of the species must be revised upwards to explain the genetic diversity observed at this locus) are probably conservative.

The results described above ignore the possibility of recurrent appearance of resistant alleles by mutation after the onset of pesticide use. Therefore, they demonstrate that the diversity observed among swept alleles can be furnished entirely by the standing variation at the time of the environmental shift. To investigate the relative importance of the standing variation versus recurrent mutation, we return to our baseline setup with *N*_*e*_ = 10^4^ and allow for a positive rate of mutation to resistant alleles after the onset of pesticide use, equal to the mutation rate beforehand. Since, when adaptation from the standing variation does not occur, recurrent mutation will eventually furnish alleles that sweep to fixation, we focus on the relative importance of the standing variation versus recurrent mutation in cases where the standing variation does supply some of the alleles involved in a selective sweep. The ‘unconditional’ importance of the standing variation versus recurrent mutation can then be calculated by simply weighting according to the probability that a sweep does occur that involves alleles from the standing variation.

When *θ* = 0.04, sweeps that involve alleles from the standing variation seldom also involve an appreciable frequency of alleles produced by recurrent mutation, especially under a dominance shift of the resistant allele (Fig. S8). However, given the rarity of sweeps that do involve alleles from the standing variation in this case (Fig. 5A), adaptation in response to pesticide use must typically exclusively involve resistant alleles produced by mutation after the onset of pesticide use.

When *θ* = 0.4, it is still the case that sweeps involving alleles from the standing variation typically make use of few or no alleles produced by recurrent mutation (Fig. S8). This is especially so under a dominance shift of the resistant allele (Fig. S8). However, in contrast to the case where *θ* = 0.04, the increased frequency of sweeps that do involve alleles from the standing variation when *θ* = 0.4 (Fig. 5A) implies that the unconditional importance of recurrent mutation for adaptation in response to pesticide use is reduced. This holds whether there is a dominance shift or not. However, it is especially strong under a dominance shift because both (i) the importance of recurrent mutation in sweeps that involve the standing variation is smaller (Fig. S8), and (ii) sweeps that do involve the standing variation are more common (Fig. 5A).

In summary: (i) The genetic diversity observed by Karasov et al. (2010) at the *Ace* locus in *D. melanogaster* can be explained with a more modestly revised estimate of the relevant effective population size of this species when the dominance shift that resistant alleles have empirically been shown to exhibit is taken into account. (ii) For the associated value of *θ*, and with a dominance shift of the resistant allele, the standing variation at the time of the environmental change is itself capable of furnishing the alleles involved in a subsequent empirical soft selective sweep. (iii) Moreover, in this case, when a soft selective sweep does occur, it is expected to predominantly involve alleles from the standing variation, rather than alleles produced by mutation after the environmental change.

## 4 Discussion

More than a century of research on the physiological and evolutionary bases of allelic dominance has been centered around, and has generated explanations for, the fact that beneficial alleles tend to be dominant while deleterious alleles tend to be recessive (Bourguet 1999; Falk 2001). Here, we have explored the implications of this pattern of fitness dominance for the genetics of adaptation to a new environment, where an allele that was deleterious in the old environment becomes beneficial in the new environment. This model is basic to the selective sweeps literature, but has typically been studied under the assumption that the fitness dominance of the focal allele is constant across the environmental change. This is contrary to the prediction of the physiological and evolutionary theories of dominance, which suggest that the allele should instead often shift from recessive deleterious before the environmental change to dominant beneficial after. We have shown that, relative to the case where the allele maintains a constant fitness dominance, the expected shift in dominance: (i) increases the probability of adaptation from the standing variation that was present at the time of the environmental change; (ii) increases the probability that multiple alleles from the standing variation will contribute to adaptation, and that these alleles will lie on distinct haplotypes; (iii) increases the probability that a soft sweep from the standing variation will be detectable in small genomic samples; (iv) increases the importance of the standing variation relative to subsequent mutation for eventual adaptation to the new environment; (v) increases the probability of evolutionary rescue when the change of environment threatens the viability of the population.

### Connections to previous theory

While most of the prior literature on selective sweeps in a new environment has assumed constant dominance of the relevant alleles, a notable exception is Orr and Betancourt (2001), who consider the case of an allele that transitions from deleterious to beneficial across an environmental change, allowing for the possibility that the allele’s fitness dominance shifts across the environmental change as well. They find that the probability that copies of the allele segregating in the standing variation at the time of the environmental change go on to fix is modulated by the ratio of the dominance of the focal allele in the new environment to its dominance in the old environment [*h*_*b*_*/h*_*d*_; Eq. 19 in Orr and Betancourt (2001)]. However, they argue that, because ‘it is hard to see why [dominance] shifts would be systematic in direction’, the dominance values of alleles across environmental shifts will tend to cancel each other out on average, and thus have no systematic effect on the probability of adaptation from the standing variation.

Orr and Betancourt’s calculations assume that, at the time of the environmental change, the allele’s copy number equals the expectation of its mutation-selection distribution from before the environmental change, although they do compare their results against numerical computations based on the full mutation-selection-drift distribution. Hermisson and Pennings (2005) carry out an analytical calculation of the probability of adaptation from the standing variation, taking into account the full mutation-selection-drift distribution of the allele. They find that, unless selection against the allele before the environmental change is weak, the probability of adaptation from the standing variation is a function of *h*_*b*_*/h*_*d*_ [Eq. 8 in Hermisson and Pennings (2005)], echoing the result of Orr and Betancourt (2001). They conclude that, when the dominance of the allele does not shift across the environmental change, its value does not matter for the probability of adaptation from the standing variation.

We have argued, based on the physiological and evolutionary theories of dominance, that systematic dominance shifts of the alleles involved in adaptation to new environments are in fact expected—these alleles are predicted often to shift from recessive when deleterious to dominant when beneficial. These expected dominance shifts facilitate adaptation from the standing variation by increasing the presence of the allele in the population prior to the environmental change, and by rescuing the allele from Haldane’s sieve after the environmental change. By incorporating the insights of the physiological and evolutionary theories of dominance into models of adaptation to new environments, we have shown that expected dominance shifts have a large impact not only on the probability of adaptation from the standing variation, but also on the genetic nature of this adaptation—in particular, whether it proceeds via hard or soft selective sweeps.

### When are dominance shifts expected?

We have outlined two scenarios under which an environmental change can cause an allele to transition from deleterious to beneficial. In the first scenario, the phenotype that is primarily under selection does not change, but the direction of selection acting on the phenotype does. Since the same phenotype is under selection across the environmental change, the fitness dominance of an allele that affects the phenotype is not expected to shift appreciably. One context where this case is expected to be especially common is domestication, where a trait that was previously suppressed by breeders might suddenly become desired [e.g., various coat properties in domestic dogs (Cadieu et al. 2009)].

In the second scenario, the change in environment corresponds to a change in the phenotype that is primarily under selection. The focal allele is deleterious before the environmental change through its association with the old phenotype under selection, and beneficial after the environmental change through its association with the new phenotype under selection. The fitness dominance of the allele is then not constrained to remain constant across the environmental change, and, indeed, theories of dominance predict that the dominance of the allele should usually increase as it transitions from deleterious to beneficial.

Environmental changes that generate selection on new phenotypes are, of course, expected to be common. We have discussed, as an example, the evolution of pesticide resistance at the *Ace* locus in insects, which encodes the enzyme acetylcholinesterase. In this case, the phenotype that was primarily under selection changed from ‘intrinsic’ enzymatic efficiency in the pesticide-free environment to the ability to inhibit pesticide binding once pesticides came into common use; accordingly, resistant alleles at *Ace* shifted from deleterious to beneficial. In agreement with the physiological and evolutionary theories of dominance, evidence suggests that resistant alleles concomitantly shifted from recessive deleterious in pesticide-free environments [based on the biochemical properties of acetylcholinesterase (Bourguet and Raymond 1998; Shi et al. 2004) and empirical measurement in moths and mosquitoes (Labbé et al. 2014; Zhang et al. 2015)] to dominant beneficial in environments of pesticide use (Bourguet and Raymond 1998; Charlesworth 1998).

In general, the arguments above suggest that adaptation to novel pesticides should be a promising arena for dominance shifts of the alleles involved; consistent with this, several further examples are already known of pesticide-resistant alleles that have undergone dominance shifts, including the alleles that confer resistance to warfarin in Norway rats (Greaves et al. 1977; Hedrick 2012) and chlorsulfuron in *Arabidopsis thaliana* (Roux et al. 2004).

A similar situation occurs when a population is exposed to new diseases or parasites. A well-known example of adaptation in humans involves the *β*-globin gene. Homozygotes for the ‘sickle-cell’ allele of this gene have characteristically misshapen red blood cells, and suffer from sickle-cell anemia, a severe blood disorder (Kwiatkowski 2005; Hedrick 2011). Heterozygotes produce functional red blood cells; the substantial deleterious effect of the allele is thus recessive (Kwiatkowski 2005). The sickle-cell allele also offers protection against malarial infection in both heterozygotes and homozygotes (Kwiatkowski 2005; Hedrick 2011), leading to a situation of heterozygote advantage in environments where the disease is prevalent (since homozygotes still suffer the severe effects of sickle-cell anemia). Thus, as for pesticide resistance, the set of phenotypes under selection changes depending on the environment: in malaria-free environments, the phenotype under selection is ‘intrinsic’ function of red blood cells, while in environments where malaria is widespread, both intrinsic function and the ability to protect against malaria are under selection.

The sickle-cell variant of *β*-globin is one of a broader class of mutations in humans—including the variants causing *α* and *β*-thalassemia, glucose-6-phosphate dehydrogenase deficiency, and cystic fibrosis—that are associated with substantial fitness costs but also confer protection against some pathogen (Clegg and Weatherall 1999; Kwiatkowski 2005; Nielsen et al. 2007; Hedrick 2012). As predicted by theories of dominance, these variants display contrasting fitness dominances in pathogen-free and pathogen-affected environments. They are recessive deleterious in pathogen-free environments, where selection acts primarily on the diseases that they cause. In environments where the pathogens that they confer resistance against are prevalent, the fitness dominance of these variants characteristically shifts all the way to overdominance. Note that this overdominance is predicted by our ‘bottom-up’ model of adaptation to a new environment, as employed in our analysis of evolutionary rescue, and is a consequence of the standard dominance patterns when there are two selectively independent phenotypes.

Environmental changes frequently expose populations to novel causes of selection (e.g., pesticides or pathogens in the examples set out above), and so often lead to shifts in the phenotypes that are primarily under selection. For this reason, shifts in the phenotypes under selection are also expected when populations colonize a new geographic area or expand into a new ecological niche. For alleles that affect both the old and new selected phenotypes, dominance shifts are expected. Thus, dominance shifts may play an important role in facilitating adaptation from the standing variation across a broad range of evolutionary and ecological contexts.

### When are dominance shifts most influential?

We have shown that dominance shifts can lead to the frequent occurrence of soft selective sweeps in parameter regimes where they ordinarily would not be expected to occur. In particular, we have shown that dominance shifts can have a very strong impact on the genetics of adaptation when the focal allele is highly deleterious prior to the environmental change (Figs. 5, S4, S9). This is because, when selection against a deleterious allele is strong, its dominance has a large effect on the shape of its mutation-selection-drift distribution, which in turn determines the likelihood of a soft selective sweep from the standing variation once the allele becomes beneficial. In contrast, if selection against the allele prior to the environmental shift is weak, there will usually be many copies present in the standing variation, regardless of its dominance, and so a soft selective sweep would be a common outcome in any case.

Our prediction that dominance shifts will be more influential when the focal allele is highly deleterious before the environmental change is complemented by a key prediction of the Wright-Kacser-Burns metabolic theory of dominance: the more deleterious a mutation is, the more recessive it usually will be (Kacser and Burns 1981; Phadnis and Fry 2005). This negative correlation between *h*_*d*_ and *s*_*d*_ has also been predicted by evolutionary theories of dominance [e.g., Manna et al. (2011)] and supported by empirical work in *Drosophila* (Simmons and Crow 1977; Charlesworth 1979) and yeast (Phadnis and Fry 2005; Agrawal and Whitlock 2011). It suggests that, precisely when dominance shifts are most influential—i.e., when the focal allele is highly deleterious before the environmental change—they should also be most pronounced (Fig. S9).

### Complex demography and selection at linked sites

The models we have studied in this paper are highly stylized, involving selection among two alleles at a single, isolated locus in a well-mixed population of constant size (this last assumption was relaxed in our analysis of evolutionary rescue). In reality, selection occurs at loci linked to any focal locus, while populations fluctuate in size over time and are structured in complex ways. All of these complications will affect the allelic frequency dynamics at a focal locus, and therefore the probability of a sweep (and of a soft sweep) in response to an environmental change.

These features vary in complex ways within genomes and across species, making it difficult to incorporate them in a general way in our model. Fortunately, to a first approximation, their influence on the adaptive process at the focal locus can be understood in terms of their effect on the relevant effective population size, *N*_*e*_, at the focal locus. Thus, purifying selection at linked sites reduces the relevant effective population size at the focal site in a way that can be captured by a single, measurable parameter, *B* (McVicker et al. 2009). Similarly, temporal changes in population size alter the effective population size in a well understood fashion (Wright 1938). Note that, as discussed above, there has recently been some interest in the timescale over which changes in population size influence the effective population size that is relevant for rapid adaptation to a new environment; the appropriate timescale will typically be much more recent than the timescale relevant for a traditional effective population size based on neutral genetic diversity (Karasov et al. 2010).

Finally, the effect of population structure depends on whether the change of environment affects the population homogenously or not. If it does, then the effect of population structure can again be understood in terms of its effect on the effective population size [e.g., (Wright 1943; Whitlock and Barton 1997)]. If not, and some subpopulations (or regions) do experience the environmental change while others do not, then the dynamics of adaptation are more complicated, with gene flow between subpopulations influencing allelic frequency dynamics and impeding adaptation in the various environments. Such a situation has occurred in the mosquito *Culex pipiens*, with spatial heterogeneity in the use of organophosphate pesticides resulting in complex geographic patterns of the frequency of pesticide-resistance mutations at the *Ace* locus (Lenormand et al. 1999; Labbé et al. 2007b).

### Implications for the genetics of adaptation across the genome

Our work has focused on the case of a single diploid locus, for which we have shown that the dominance shifts predicted by the evolutionary and physiological theories of dominance lead to an increased probability of adaptation to a new environment from the standing variation, and an increased probability that multiple alleles will be involved in this adaptation—i.e., an increased probability of a soft selective sweep. This result relies on the possibility of strong fitness dominance, and therefore points to differences across the genome in where hard versus soft sweeps are likely to occur.

First, while autosomal loci are diploid in both sexes, and therefore subject to the patterns of the genetics of adaptation described in this paper, X-linked loci in male-heterogametic systems and Z-linked loci in female-heterogametic systems are diploid in one sex but haploid in the other. At these sex-linked loci, alleles that become beneficial in a new environment cannot have been strongly recessive deleterious in the old environment, because they are hemizygously expressed in one of the sexes. Therefore, we predict that soft sweeps from the standing variation should be less common at sex-linked loci than at autosomal loci. This prediction is, of course, complicated by other differences between autosomes and sex chromosomes, such as differences in their effective population size and the strength of selection on males versus females (Vicoso and Charlesworth 2006).

Second, at a single diploid locus, the fitness effects of a deleterious allele can be ‘masked’ by the wild-type allele at the same locus, but when there are two (or more) copies of a locus, a deleterious allele can be masked by a wild-type allele at its locus or at the other locus. Therefore, we expect a deleterious allele at a duplicated locus to be more recessive than it would be in the single-locus case, since it has potentially more wild-type alleles to mask its deleterious effect. If the allele were later to become dominant beneficial, a soft sweep would then be an even more likely outcome than in the single-locus case, because the allele would undergo a more extreme dominance shift. This increased likelihood of a soft sweep would be further enhanced by the larger mutational target presented by the duplicated locus, i.e., a higher effective value of *θ*.

Note that, if, after the change of selective environment, a sweep does occur at a duplicated locus, this would appear as neofunctionalization of the gene (Ohno 1970; Force et al. 1999). However, another source of apparent neofunctionalization is suggested by the the ‘bottom up’ model of dominance shifts that we used to study the particular case of evolutionary rescue. Under this model, heterozygote advantage is expected to be a common outcome of changes in the selective environment, leading to a partial selective sweep and subsequent stable polymorphism at the focal locus. In a sexual species, Mendelian segregation at the polymorphic locus causes the production of less fit homozygous genotypes, inducing a ‘segregation load’ that can select for duplication of the locus and fixation of the alternative alleles, one at each locus (Haldane 1954; Spofford 1969; Milesi et al. 2017). These two sources of neofunctionalization could be empirically distinguished if the timing of the duplication were either known *a priori* or inferable through the sequence divergence of the two gene copies. A case where the chronology of gene duplication is well established is the *Ace* locus in *Culex pipiens* (Labbé et al. 2007a). In multiple populations subjected to pesticides, haplotypes that harbor a duplication of the *Ace* locus, with a copy each of the susceptible and resistant allele, are undergoing selective sweeps (Labbé et al. 2007a; Alout et al. 2011; Milesi et al. 2017). The duplication is known to have occurred after the onset of pesticide use (Labbé et al. 2007a), consistent with the second scenario for neofunctionalization outlined above.

## 5 Conclusion

We have shown that dominance shifts have a major impact on the genetics of adaptation to a new environment, increasing the likelihood of selective sweeps and of soft selective sweeps in particular. To the extent that dominance shifts in response to a change in environment are common—as physiological and evolutionary theories of dominance predict they should be—our findings clearly have important implications for the genetic patterns that will be observed following adaptation. Unfortunately, although there have been many cases where the alleles involved in adaptation to new environments have been identified, it is only in a handful of these cases that the dominances of the allele before and after the environmental change have been measured. In showing that dominance shifts (i) are expected to be common and (ii) can have a major impact on the genetics of adaptation, we hope that our results will encourage geneticists interested in adaptation to new environments to measure dominance in investigations of alleles that have undergone selective sweeps.

## Acknowledgments

We are grateful to Graham Coop, Nate Edelman, Joachim Hermisson, Thomas Lenormand, Pleuni Pennings, Peter Wilton, and members of the Coop lab at UC Davis for helpful discussions, and to Graham Coop and the referees for comments that improved the manuscript. PM is supported by a Center for Population Biology postdoctoral fellowship and an NSF postdoctoral fellowship. CV is supported by a Branco Weiss fellowship. This work was supported in part by the National Institute of General Medical Sciences of the National Institutes of Health (grants NIH R01 GM108779 and R35 GM136290 to G. Coop). The computations in this paper were run on the FASRC Odyssey cluster supported by the FAS Division of Science Research Computing Group at Harvard University.

## Code availability

Simulation code is available at https://github.com/Pavitra451/dominanceshifts.

---

**Figure S1:**
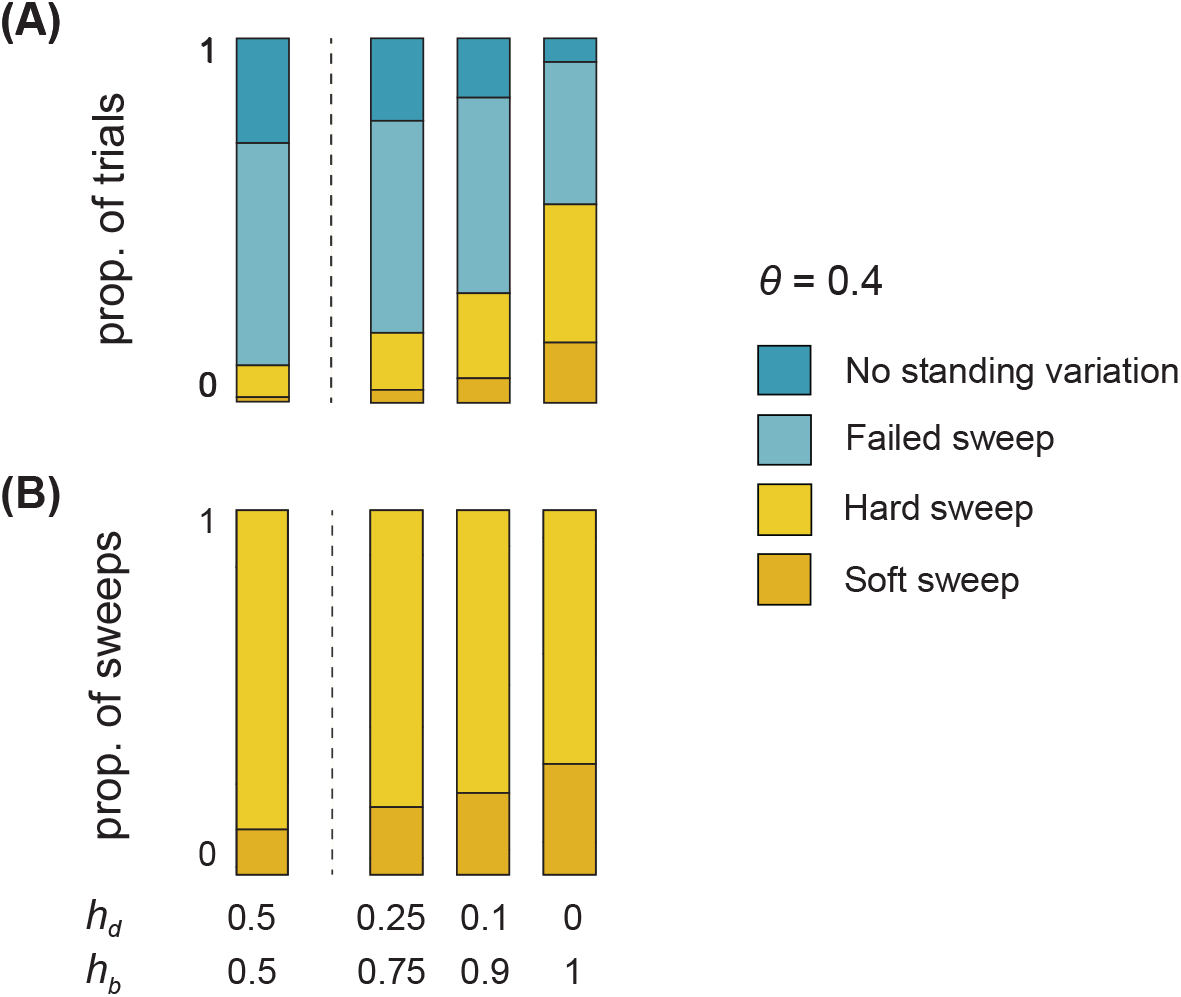
Dominance shifts increase (A) the likelihood of a selective sweep from the standing variation, and (B) the relative likelihood of soft versus hard selective sweeps, both when the beneficial and deleterious selection coefficients of the focal allele are fixed (Figs. 1, S4) and when they are random variables drawn from empirically justified distributions of fitness effects (this figure).

**Figure S2:**
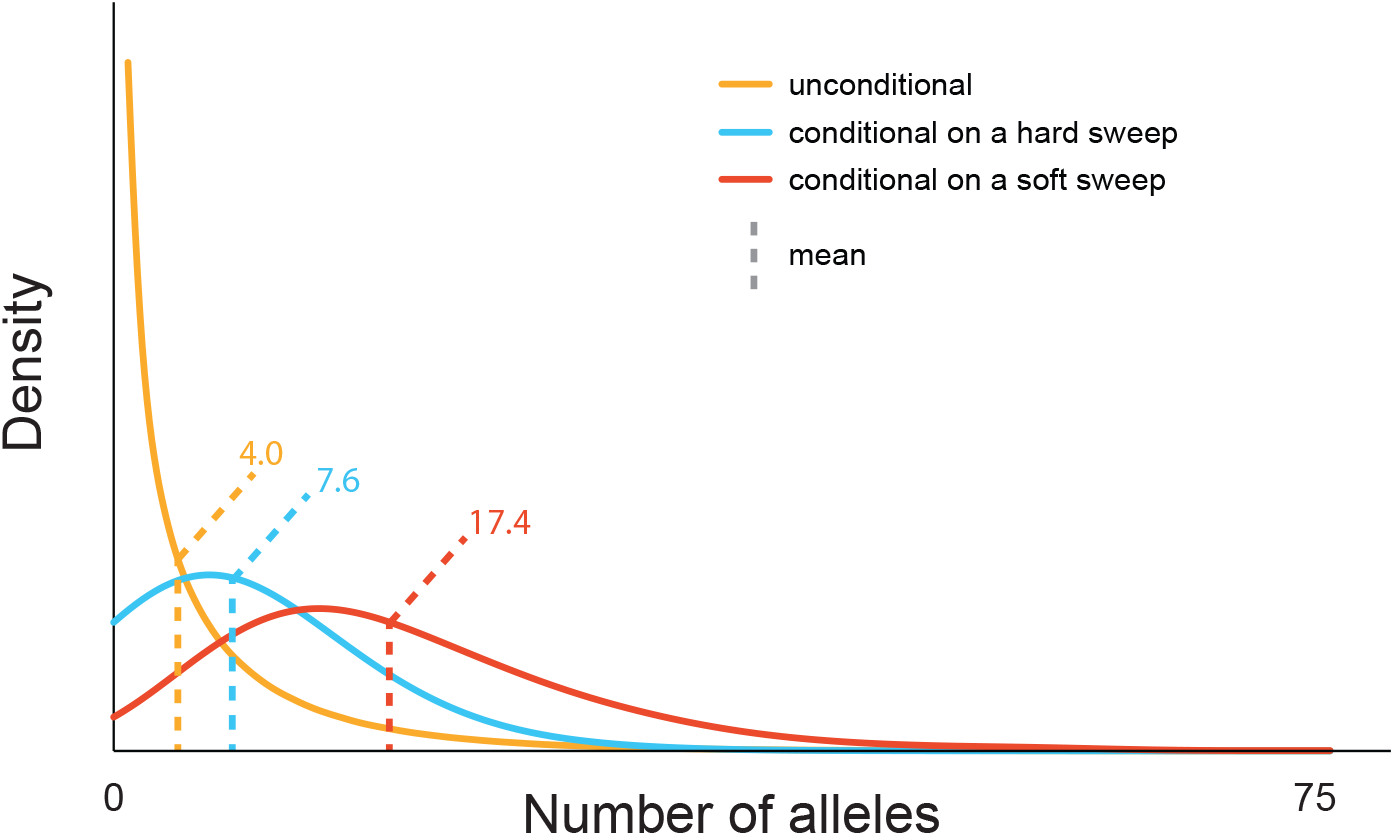
Hard and soft selective sweeps derive from different regions of the mutation-selection-drift distribution. The mutation-selection-drift distribution of a deleterious allele is shown in gold [*θ* = 0.4 (*N* = 10^4^, *u* = 10^−5^), *s*_*d*_ = 0.1, *h*_*d*_ = 0.5], while the distributions of starting allele counts that result in hard (blue) and soft (red) selective sweeps are overlaid (*s*_*b*_ = 0.1, *h*_*b*_ = 0.5).

**Figure S3:**
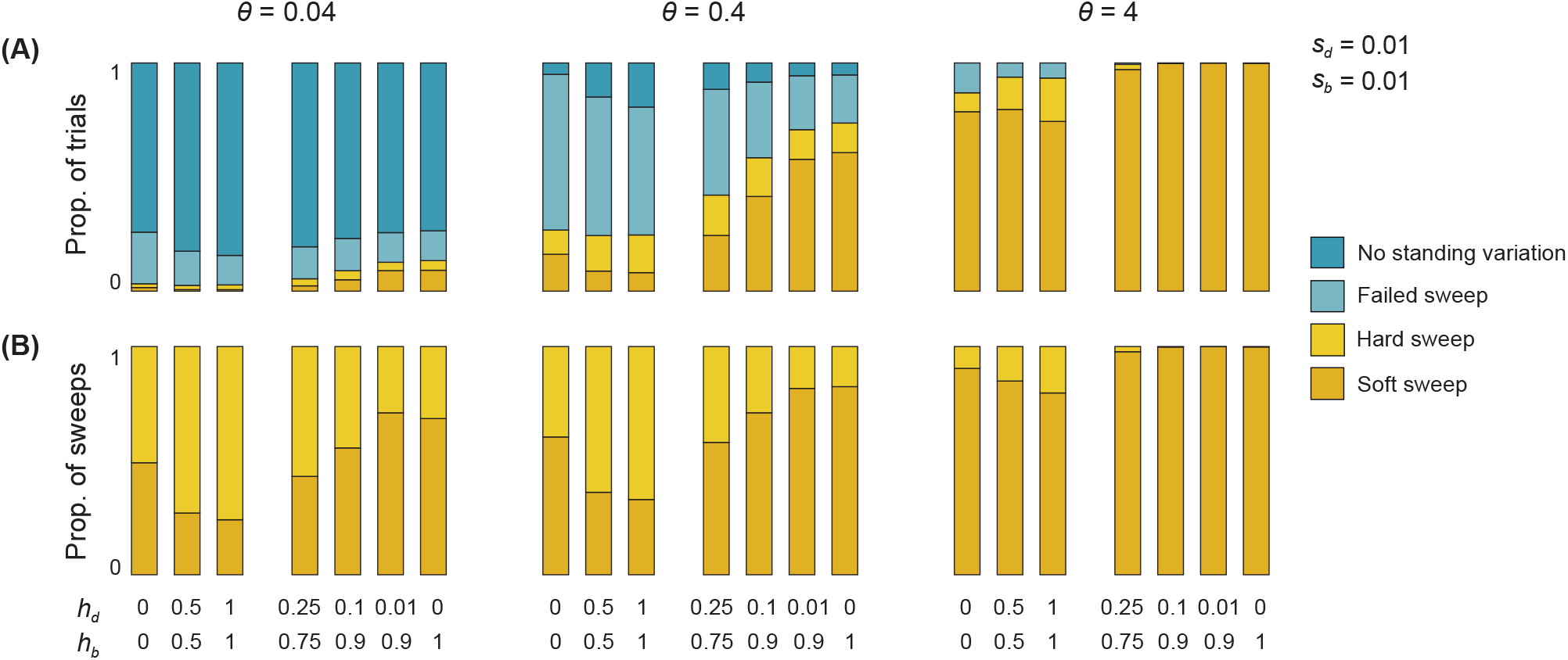
(A) Expected dominance shifts increase the likelihood of a selective sweep relative to scenarios in which the adaptive allele is constantly recessive, additive, or dominant. (B) Conditional on successful adaptation from the standing variation, expected dominance shifts increase the probability that multiple alleles are involved. When the allele maintains a constant dominance across the environmental shift (*h*_*b*_ = *h*_*d*_), the relative likelihood of soft sweeps is highest when it is fully recessive (*h*_*d*_ = *h*_*b*_ = 0). Compared to this case, a moderate dominance shift (e.g., *h*_*d*_ = 0.25, *h*_*b*_ = 0.75) leads to approximately the same relative likelihood of a soft selective sweep (and stronger dominance shifts increase this relative likelihood further). Moreover, since the probability of adaptation from the standing variation is lower for a constantly recessive allele than for an allele undergoing a dominance shift (A), there will be a greater absolute number of soft sweeps in the latter case.

**Figure S4:**
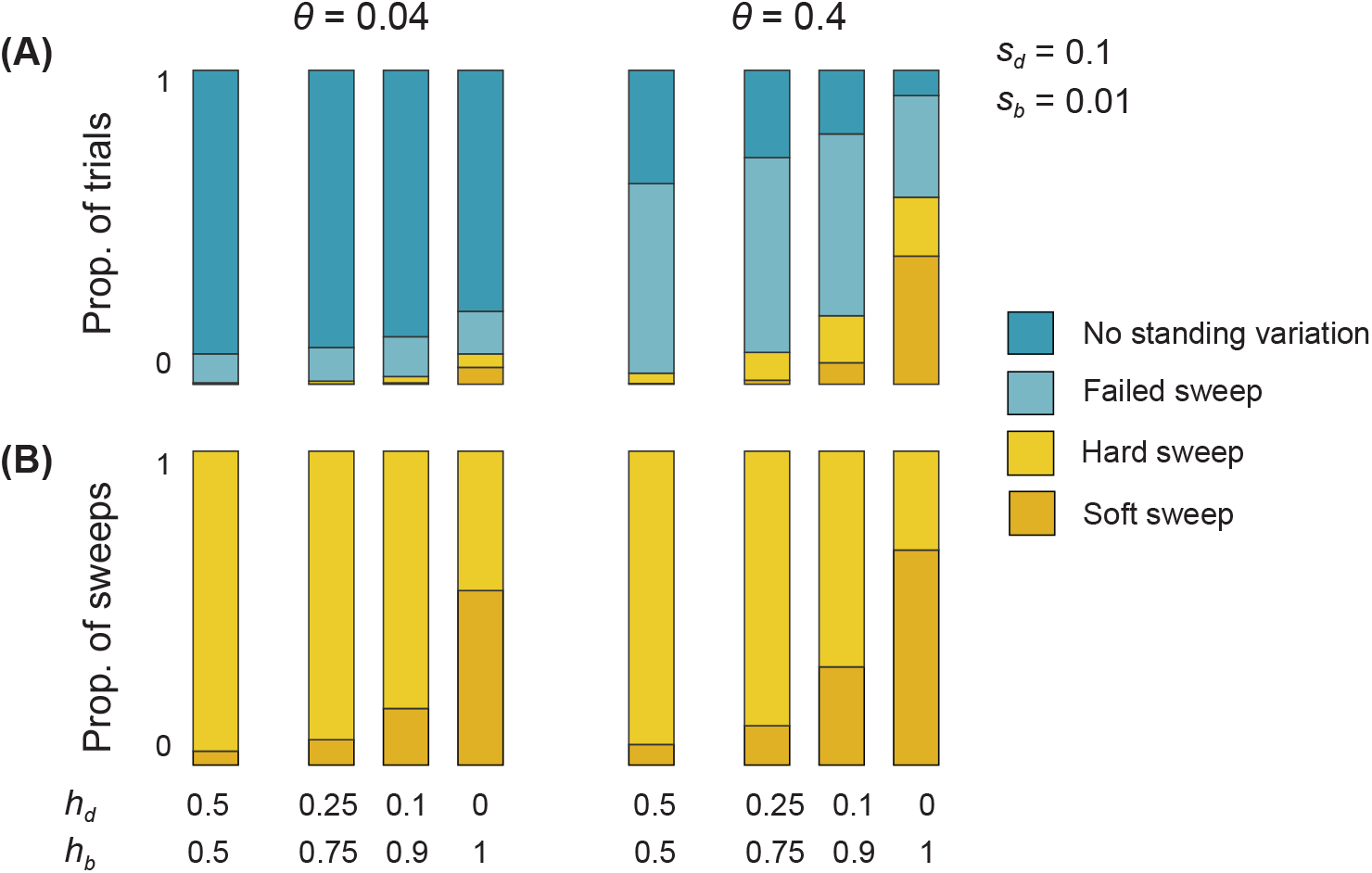
Expected dominance shifts can cause soft sweeps to predominate over hard sweeps even when *s*_*b*_*/s*_*d*_ is small. Previous work predicts that soft sweeps should be rare when *s*_*b*_*/s*_*d*_ < 1, owing to low levels of standing variation at the time of the environmental change and a high probability of subsequent stochastic loss of those copies that are present. However, if the focal allele undergoes a strong dominance shift (*h*_*d*_ ≈ 0, *h*_*b*_ ≈ 1), soft sweeps can be more likely than hard sweeps despite small values of *s*_*b*_*/s*_*d*_.

**Figure S5:**
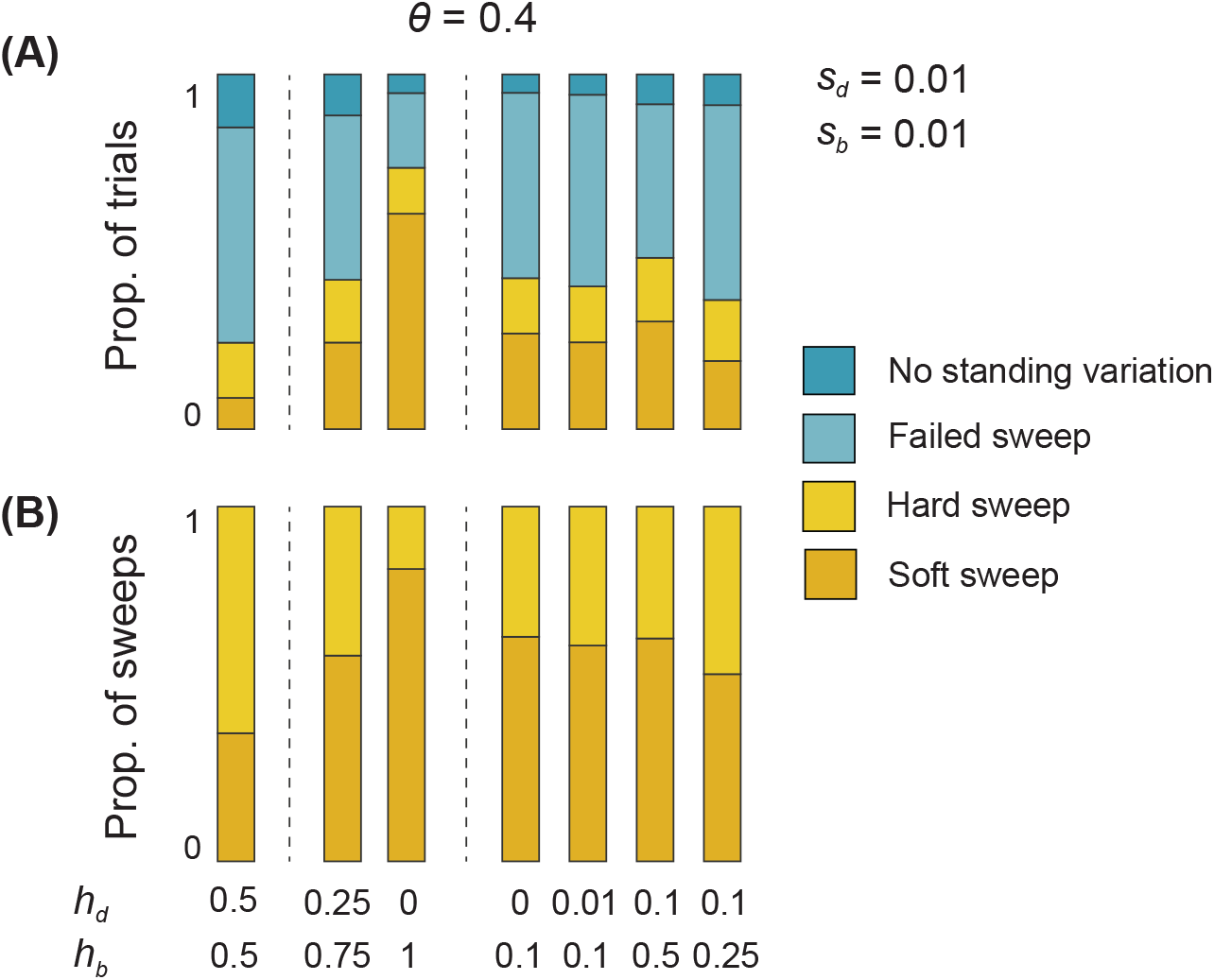
Expected dominance shifts substantially increase the likelihood of soft selective sweeps, even when the dominance of the focal allele after the environmental change is sub-additive (*h*_*d*_ < *h*_*b*_ < 0.5). However, such partial dominance shifts lead to smaller increases in the likelihood of a soft sweep, relative to complete dominance shifts (*h*_*d*_ ≈ 0, *h*_*b*_ ≈ 1).

**Figure S6:**
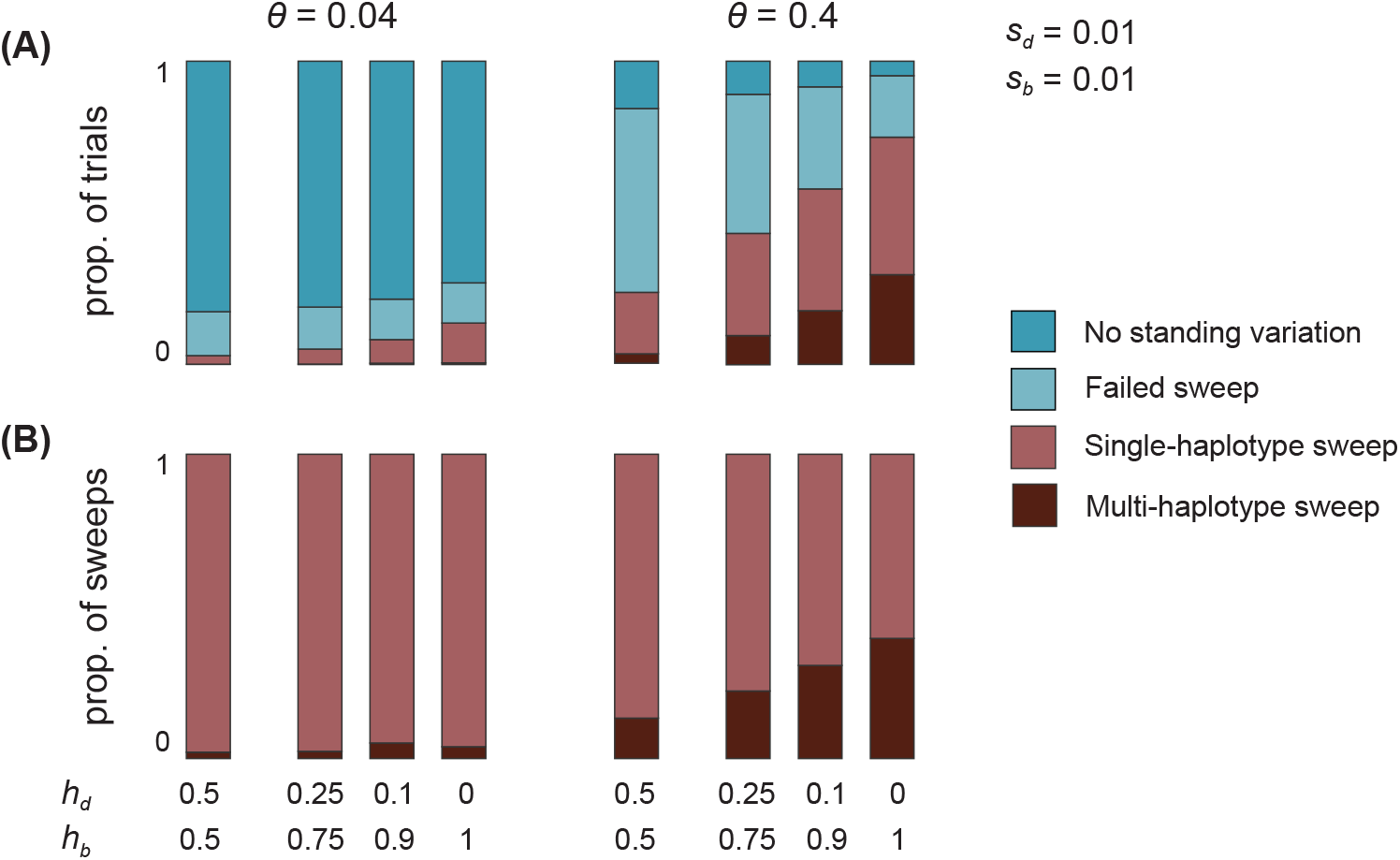
Expected dominance shifts increase the probability of soft sweeps with multiple independent mutational origins, and therefore multiple distinguishable haplotypes (soft sweeps by the ‘sample definition’).

**Figure S7:**
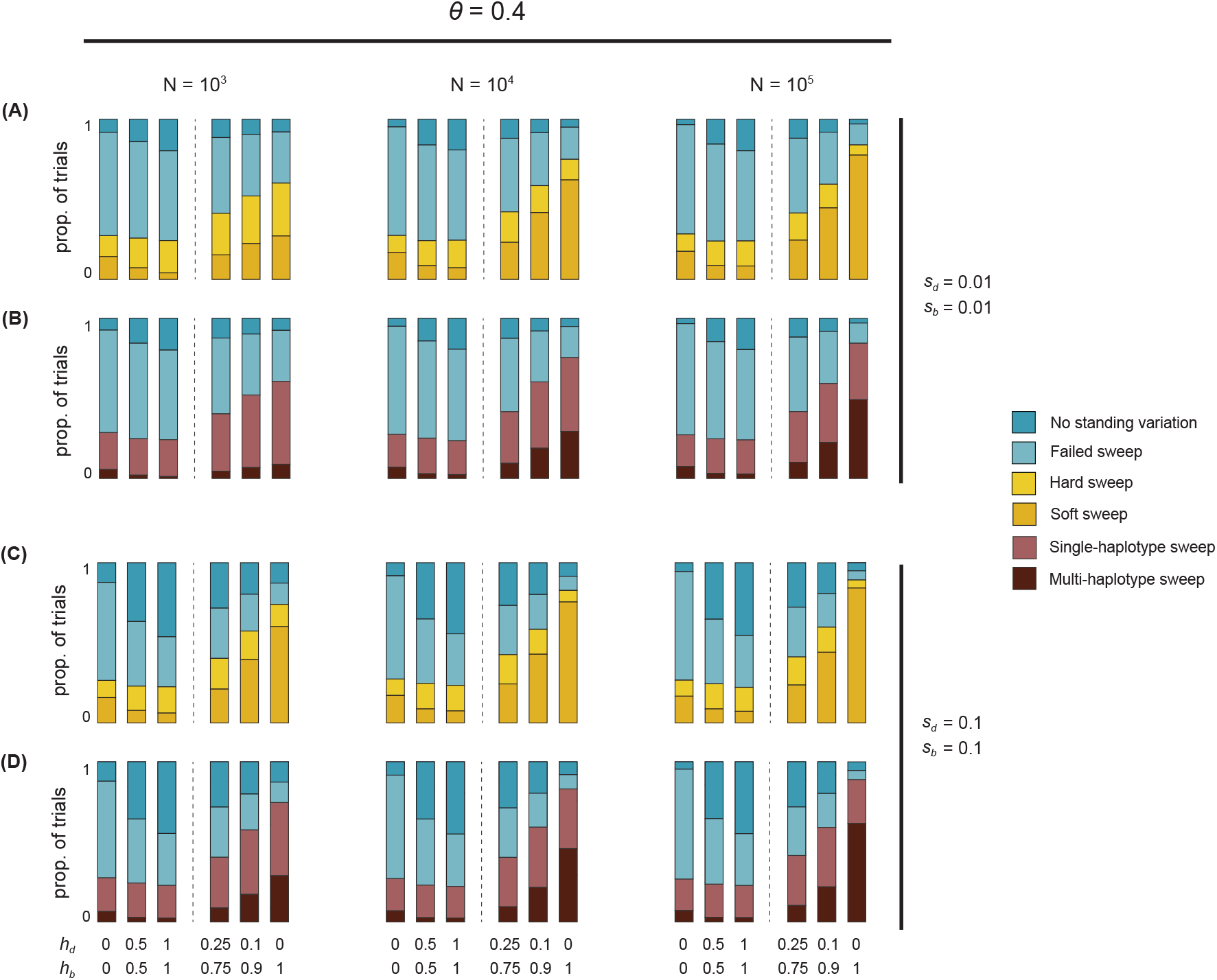
Holding the rate of mutational supply (*θ* = 4*Nu*) fixed, an increase in the effective population size *N* (and a concomitant decrease in the mutation rate *u*) does not substantially affect the probability of a sweep and the relative likelihoods of soft versus hard sweeps when the focal allele does not undergo a dominance shift (*h*_*d*_ = *h*_*b*_), but does increase the probability of a sweep and the relative likelihood of soft sweeps when the focal allele undergoes a dominance shift, especially when it undergoes a full dominance shift (*h*_*d*_ = 0, *h*_*b*_ = 1). These results hold for both the population-based definition of a soft sweep (A,C) and the sample-based definition (B,D). Panels C and D employ parameters relevant to the case of adaptation at the *Ace* locus in *Drosophila melanogaster*.

**Figure S8:**
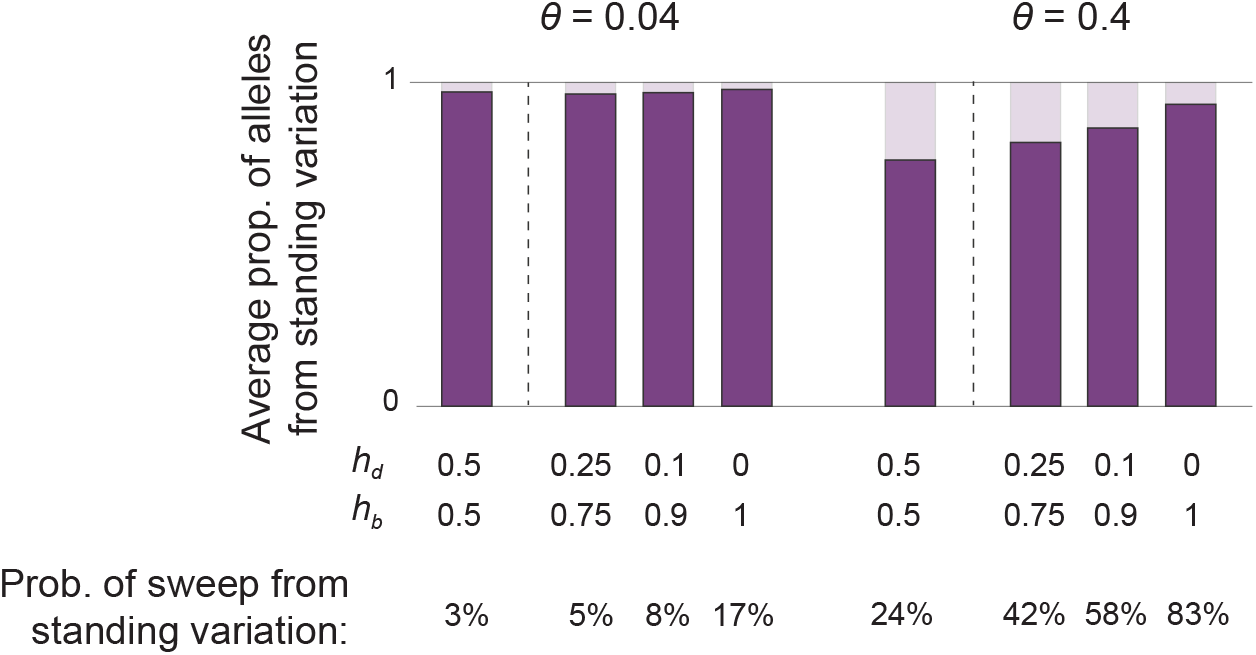
A dominance shift of resistant alleles at the *Ace* locus in *Drosophila melanogaster* after the onset of pesticide use increases the average representation of the standing variation, relative to recurrent mutation, among successful sweeps that involve some alleles from the standing variation. *θ* values correspond to the long-term effective population size of *D. melanogaster* (left) and an increased value based on more recent demography of the species (right), following the logic of Karasov et al. (2010). For the smaller value of *θ*, if a sweep occurs that involves alleles from the standing variation (although it seldom does; see bottom panel), it is expected to be dominated by alleles from the standing variation, regardless of whether a dominance shift occurs or not. In contrast, for the larger value of *θ*, dominance shifts substantially increase the representation of standing variation among sweeps that involve some alleles from the standing variation (which also become common).

**Figure S9:**
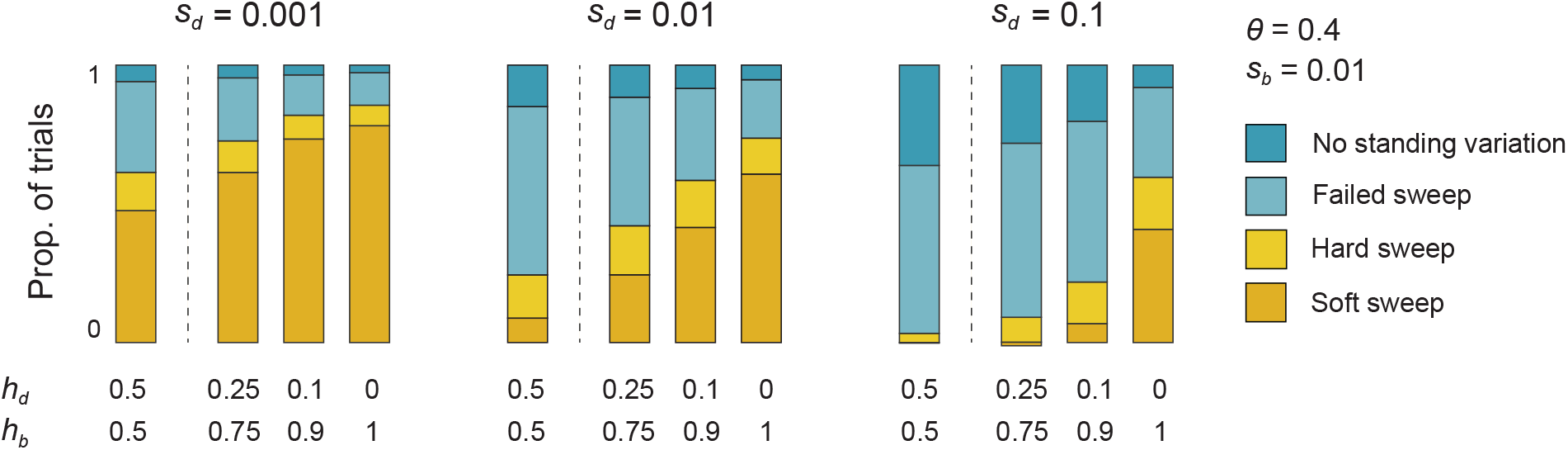
Expected dominance shifts have the largest proportional effect on the probability of adaptation from the standing variation, and the relative proportion of soft versus hard sweeps, when selection against the focal allele prior to the environmental change is strong. Complementing this effect, theories of dominance predict a negative correlation between deleterious effect size and dominance, such that more deleterious alleles are typically more recessive. Thus, dominance shifts are expected to play a particularly important role in parameter regimes in which the focal allele is strongly selected against before the environmental change.

**Figure S10:**
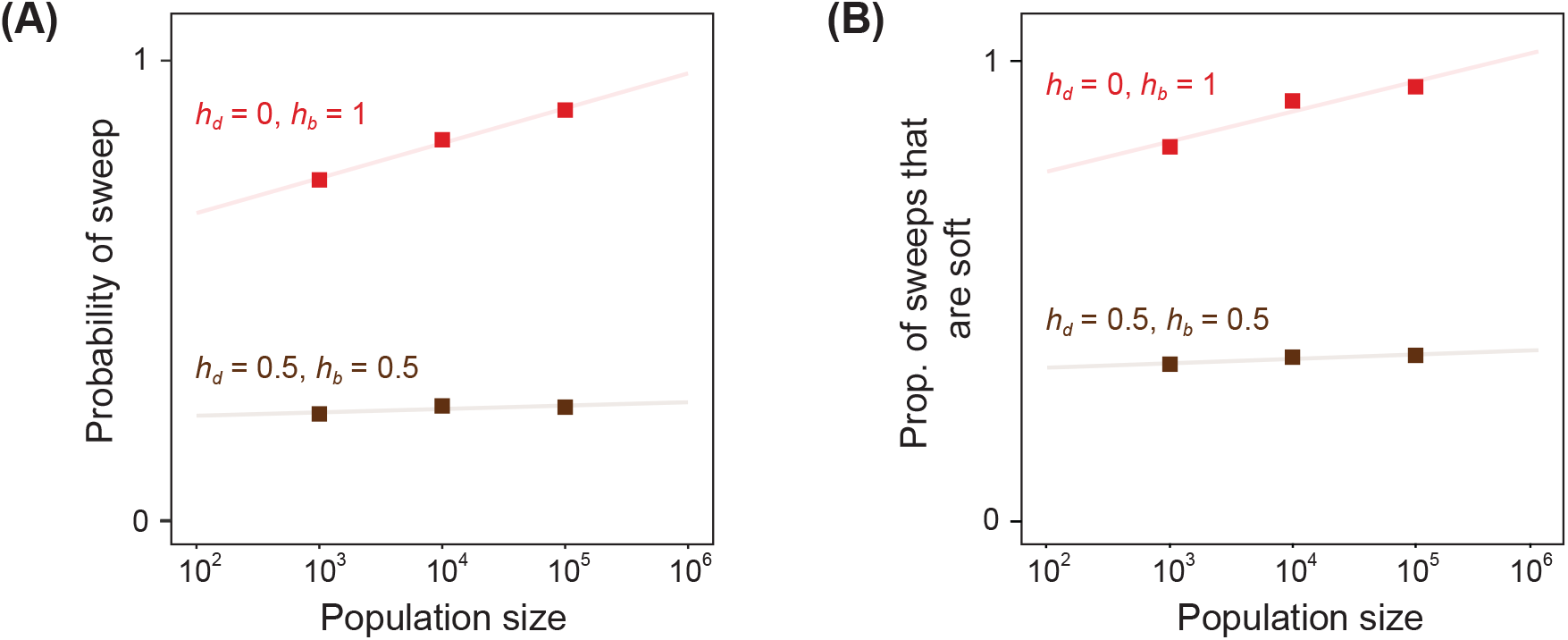
Holding *θ* = 4*Nu* constant, increasing the effective population size increases the likelihood of selective sweeps (A), and soft selective sweeps relative to hard selective sweeps (B), only when there is a dominance shift of the focal allele. The results displayed here are isolated from Fig. S7, in order to permit rough extrapolation to larger effective population sizes, including effective population sizes relevant to adaptation in *Drosophila melanogaster* (*N*_*e*_ ≥ 10^6^). Parameters are chosen to match adaptation at the *Ace* locus in *D. melanogaster*: *s*_*d*_ = 0.1 and *s*_*b*_ = 0.1.

